# Variation in subcortical anatomy: relating interspecies differences, heritability, and brain-behavior relationships

**DOI:** 10.1101/2022.04.11.487874

**Authors:** Nadia Blostein, Gabriel A. Devenyi, Sejal Patel, Raihaan Patel, Stephanie Tullo, Eric Plitman, Manuela Costantino, Ross Markello, Olivier Parent, Saashi A. Bedford, Chet C. Sherwood, William D Hopkins, Jakob Seidlitz, Armin Raznahan, M. Mallar Chakravarty

**Author notes:** Correspondence (Twitter: BlosteinNadia) and (Twitter: mallarchkrvrty1).

## Abstract

There has been an immense research focus on the topic of cortical reorganization in human evolution, but much less is known regarding the reorganization of subcortical circuits which are intimate working partners of the cortex. Here, by combining advanced image analysis techniques with comparative neuroimaging data, we systematically map organizational differences in striatal, pallidal and thalamic anatomy between humans and chimpanzees. We relate interspecies differences, a proxy for evolutionary changes, to genetics and behavioral correlates. We show that highly heritable morphological measures are significantly expanded across species, in contrast to previous findings in the cortex. The identified morphological-cognitive latent variables were associated with striatal expansion, and affective latent variables were associated with more evolutionarily-conserved areas in the thalamus and globus pallidus. These findings provide new insight into the architecture of these subcortical hubs and can provide greater information on the role of these structures in health and illness.

## 1. Introduction

The basal ganglia and thalamic structures are critically involved in many cortico-subcortical circuits: the striatum receives a variety of excitatory signals from the cerebral cortex, which are modulated by the pallidum and relayed back to more specific cortical areas through the thalamus (Middleton and Strick, 2000; Grahn, Parkinson and Owen, 2008). Neurodegenerative signatures related to motor disorders such as Parkinson’s disease (Kish, Shannak and Hornykiewicz, 1988; Owens-Walton *et al*., 2018) and Huntington’s disease (Aylward *et al*., 1996; Majid *et al*., 2011) have led to a significant understanding of circuitry involving the striatum, thalamus and globus pallidus (Herrero, Barcia and Navarro, 2002). While the focus of many investigations involving subcortical structures has been related to motor performance (Brooks, 1995; Groenewegen, 2003; Lehéricy *et al*., 2006; Turner and Desmurget, 2010) there is a rapidly expanding body of literature on the role of these structures in various neuropsychiatric disorders (Konick and Friedman, 2001; de Jong *et al*., 2008; Goldman *et al*., 2008; Koolschijn *et al*., 2009; Ebdrup *et al*., 2010; Haller, 2011; Dougherty *et al*., 2012; Shepherd *et al*., 2012; Haijma *et al*., 2013; Bootsman *et al*., 2015; Chakravarty *et al*., 2015; Roalf *et al*., 2015; Schuetze *et al*., 2016; Kälin *et al*., 2017; Perani *et al*., 2018) and as key network hubs that support largely human-specific (Tranel, Cooper and Rodnitzky, 2003) cognitive behaviors (van Schouwenburg, den Ouden and Cools, 2010; Saalmann and Kastner, 2015; Anderson *et al*., 2018; Robert *et al*., 2021), often defined as *higher-order behaviors*. The role of these structures is therefore not constricted to evolutionarily conserved motor and affective behaviors, typically defined as *lower-order behaviors*. Given the role of the basal ganglia and thalamus as key structures involved in, but not limited to, two motor circuits (Alexander, DeLong and Strick, 1986), two circuits involving the prefrontal cortex (Alexander, DeLong and Strick, 1986) and a limbic circuit (Alexander, DeLong and Strick, 1986), understanding the factors that may impact their architecture is critically important.

Existing cortical studies provide a foundation for examining these factors. These studies have identified homologies in the topological patterning of cortical morphology as they relate to heritability and evolutionary expansion indexed by comparisons of nonhuman primates (such as the rhesus macaque) to humans (Patel *et al*., 2018; Wei *et al*., 2019; Valk *et al*., 2020; Jeong *et al*., 2021). For instance, surface area measurements in unimodal areas of the primary visual cortex and dorsal visual stream (Hebart and Hesselmann, 2012) are highly heritable (Patel *et al*., 2018) and highly conserved between humans and the macaque (Reardon *et al*., 2018). On the other hand, the rostrolateral and dorsolateral prefrontal cortices, involved in higher-order cognitive functions such as memory and attention (Kane and Engle, 2002), are strongly expanded across species (Reardon *et al*., 2018) and only moderately heritable (Patel *et al*., 2018), suggesting that a large portion of their variation is influenced by environmental factors. Regions that demonstrate limited interspecies differences between humans and nonhuman primates are often considered as being evolutionarily conserved. Conserved neuroanatomical regions that are influenced by genetics (highly heritable) are thought to mediate behaviors shared across species. The inverse interpretation is ascribed to cortical regions that are human-expanded and more influenced by environmental factors. Regions exhibiting this patterning are often considered to be involved in cortical adaptation to human-specific needs requiring brain plasticity (Chaplin *et al*., 2013; Sherwood and Gómez-Robles, 2017; Sneve *et al*., 2019). Such cortical areas have further been shown to be less heritable in humans than in chimpanzees (Gómez-Robles *et al*., 2015). Importantly, these observations may have significant implications in understanding human health and disease. Regions significantly expanded in the human cortex relative to nonhuman primates (Reardon *et al*., 2018) disproportionately represent areas vulnerable to neurodegenerative disorders such as Alzheimer’s Disease (Jacobs *et al*., 2012; Gefen *et al*., 2018; Prawiroharjo *et al*., 2020) and other forms of dementia (Barber *et al*., 2001; Chan *et al*., 2001; Corbett *et al*., 2009). In fact, there is evidence from genetics studies that suggests that humans evolved elements involving DNA sequence changes that converge on cell types and laminae involved in human cortical expansion and risk genes implicated in maladaptive neurodevelopment (Won *et al*., 2019). Taken together, these previous studies motivate further neuroanatomical subcortical studies with the closest living relatives of our species, the chimpanzee and other great apes.

We have limited information on these homologous properties in the subcortex: namely the fine-grained examination of genetic effects on structure, their relationship to higher-order cognition, and the relevance of these two variables in the context of interspecies differences as a means of inferring their relationship to evolutionary changes. Given the complex underlying cellular and structural properties of the striatum (Tian *et al*., 2010; Tepper *et al*., 2018), thalamus (Torrico and Munakomi, 2020) and globus pallidus (François *et al*., 1984; Yelnik, Percheron and François, 1984; Neudorfer *et al*., 2020), it is critical to leverage computational tools to characterize these structures in a fine-grained and topologically-specific manner. To this end, several studies from our group, and others, have demonstrated that surface-based morphometry measures can be used to describe local neuroanatomical phenotypes that may be missed when considering classical volumes (Kim *et al*., 2012; Raznahan *et al*., 2014; Sandman *et al*., 2014; Voineskos *et al*., 2015; Reardon *et al*., 2018; Tullo *et al*., 2019; Bussy *et al*., 2020). In the current study, we examine the patterning of human-chimpanzee interspecies differences as well human interindividual differences in subcortical anatomy and explored how this relates to other axes of anatomical variation. Specifically, we investigated how interspecies subcortical shape differences relate to human patterns of heritability (how variation in a trait is attributable to genetic effects) using a twin and non twin sibling design. Next, within humans, we test for relationships between morphological variation in the subcortex and variation in a large battery of cognitive and behavioral measures. Elucidating the interaction between genetics (heritability), environment, behavior and subcortical neuroanatomy is crucial to deepen our understanding of how the striatum, thalamus and globus pallidus are involved in normative brain function and neuropsychiatric disease.

## 2. Results

To study the spatial concordance between interspecies, intraspecies and behavior-defined influences on subcortical (striatum, thalamus, globus pallidus) patterning, we used two different datasets. In the context of this work, the terms “pallidum” and “pallidal” will be referring exclusively to the globus pallidus (i.e. the dorsal pallidum). First, interspecies differences between subcortical surface-based measures in a human structural magnetic resonance imaging (MRI) template and a chimpanzee MRI template were used to identify areal (surface area) and surface displacement properties inferred to be related to evolutionary divergence between chimpanzees and humans (see Figure 1.1; STAR Methods 3.1 and 4.1). Next, topologically-specific heritability estimates were derived from the MRI data of 1113 healthy young adults from the Human Connectome Project (HCP; S1200 Release) using a twin and non twin sibling design (see Figure 1.2; STAR Methods 3.2 and 4.2). Surface-based measures were then related to behavior in the HCP using multivariate techniques (partial least squares correlation [PLSC] analysis) (see Figure 1.3; STAR Methods 3.2 and 4.3). Finally, to identify potential overlaps between our 6 sets (3 analysis types x 2 shape measures) of subcortical brain maps, we adapted the (Alexander-Bloch *et al*., 2018) surface-based cortical spin test to subcortical structures (see Figure 1.4; STAR Methods 4.4) to derive correlations between maps and assess their significance.

**Figure 1:**
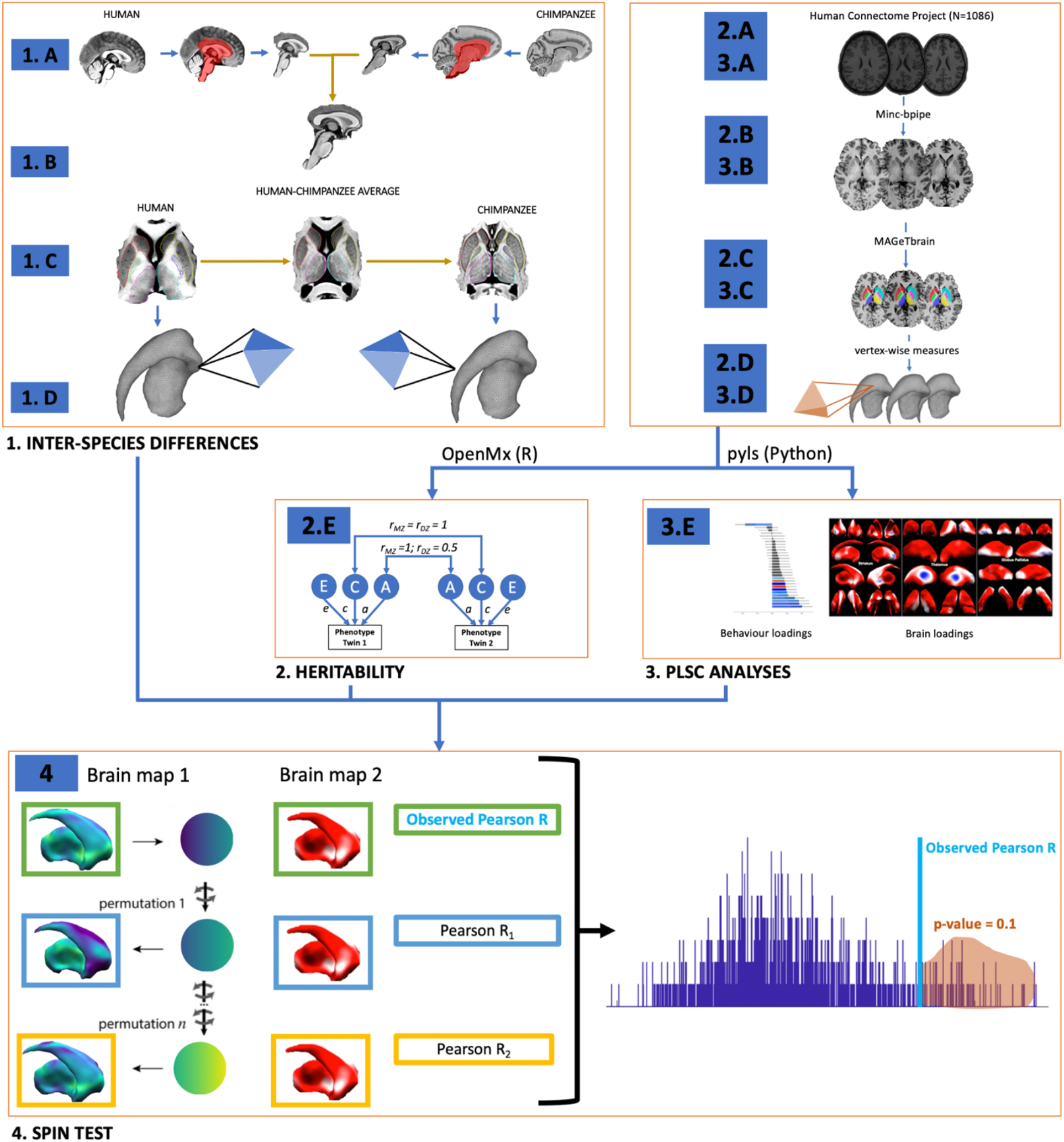
Workflow diagram. **1)** Data processing workflow for interspecies differences analysis. **2/3)** HCP data processing workflow (2/3 A-D) for heritability analysis (2 E) and PLSC analysis (3 E). **4)** Adaptation of cortical surface spin test (Alexander-Bloch *et al*., 2018) to the striatum, thalamus and globus pallidus to examine the significance of cross-analysis pairwise combinations of brain maps.

### 2.1 Interspecies differences

To make inferences about neuroanatomical evolution, we investigated interspecies differences between humans and chimpanzees. The subcortical structure surfaces from the human MAGeT brain atlases (Winterburn *et al*., 2013; Park *et al*., 2014; Tullo *et al*., 2018) were warped to fit the subcortical anatomy of the chimpanzee template, using the nonlinear transformations generated to estimate human-chimpanzee hybrid subcortical space (see Figure 1.1). We then investigated the human-to-chimpanzee ratio of vertex-wise surface area as a proxy for relative expansion or conservation in humans to chimpanzees and the vertex-wise displacement mapping from chimpanzees to humans.

#### 2.1.1 Volumes

Total brain volume (TBV) as well as ipsilateral structure-specific volumes in the striatum, thalamus and globus pallidus are higher in humans relative to chimpanzees. The volumetric expansion in humans compared to chimpanzees is noticeably greater in TBV (ratio of 3.71) relative to ipsilateral subcortical structure volumes (ratios ranging from 1.52 in the right globus pallidus to 2.13 in the left thalamus) (see Figure 2).

**Figure 2.**
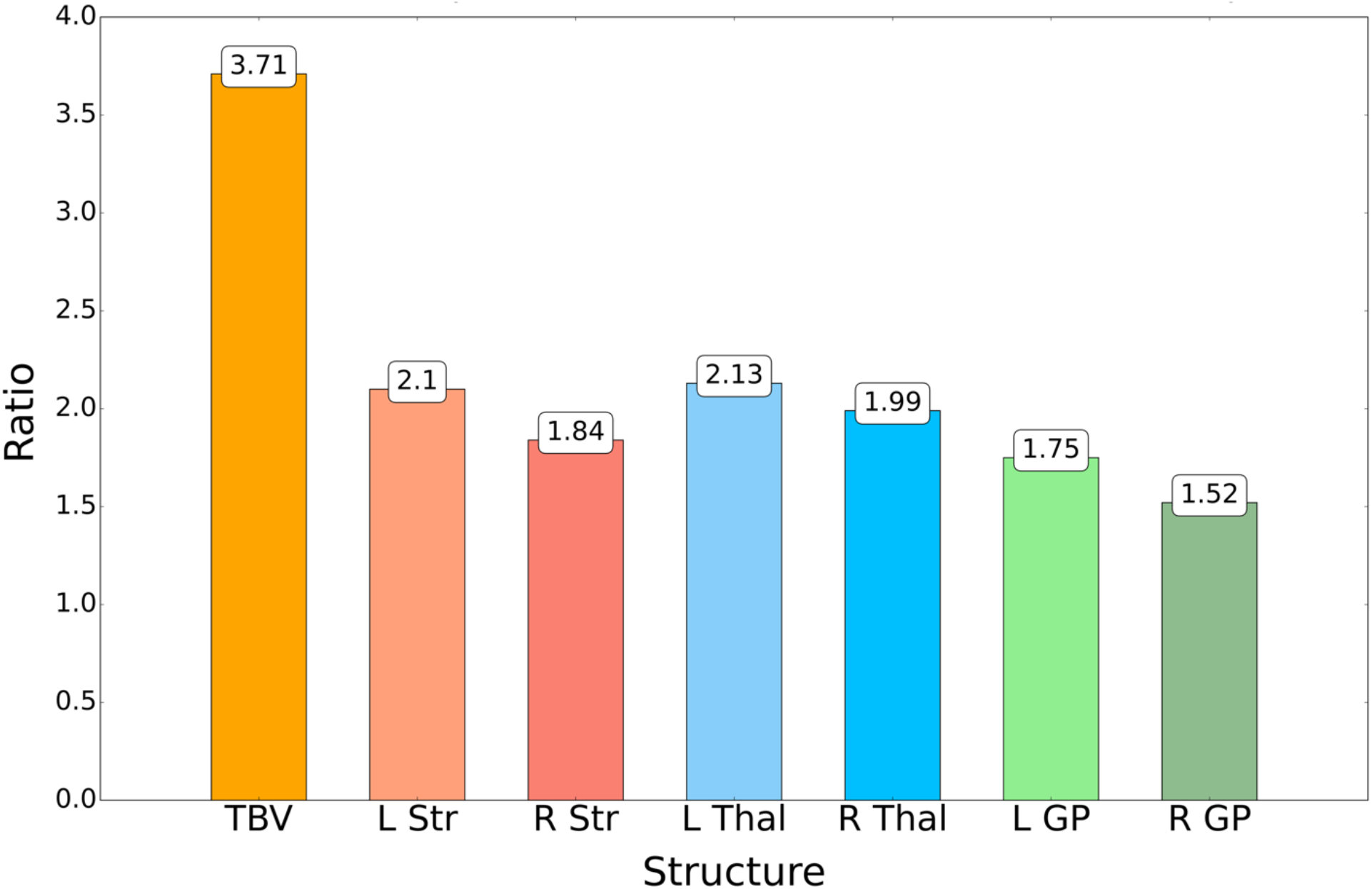
Ratios of structure-specific volumes in humans to structure-specific volumes in chimpanzees. Shown for TBV and the left (L) and right (R) striatum (Str), thalamus (Thal) and globus pallidus (GP).

#### 2.1.2 Vertex-wise measures

Before computing vertex-wise measurements on species-specific surfaces, the surfaces were adjusted for interspecies scaling differences to only examine proportional nonlinear differences. This ensures that vertex-wise measurements capture mappings that are independent of global linear effects, such as translation (see STAR Methods 3.1). Given that the subcortical volumes in humans are higher than in chimpanzees, areas where the areal ratio is above 1 or where the outward displacement from chimpanzees to humans is positive are referred to as being more expanded in humans relative to chimpanzees (red; see Figure 3). On the other hand, areas where the ratio of vertex-wise surface area in humans to chimpanzees is below 1 or where the outward displacement from chimpanzees to humans is negative are referred to as being more conserved across species (blue; see Figure 3).

**Figure 3.**
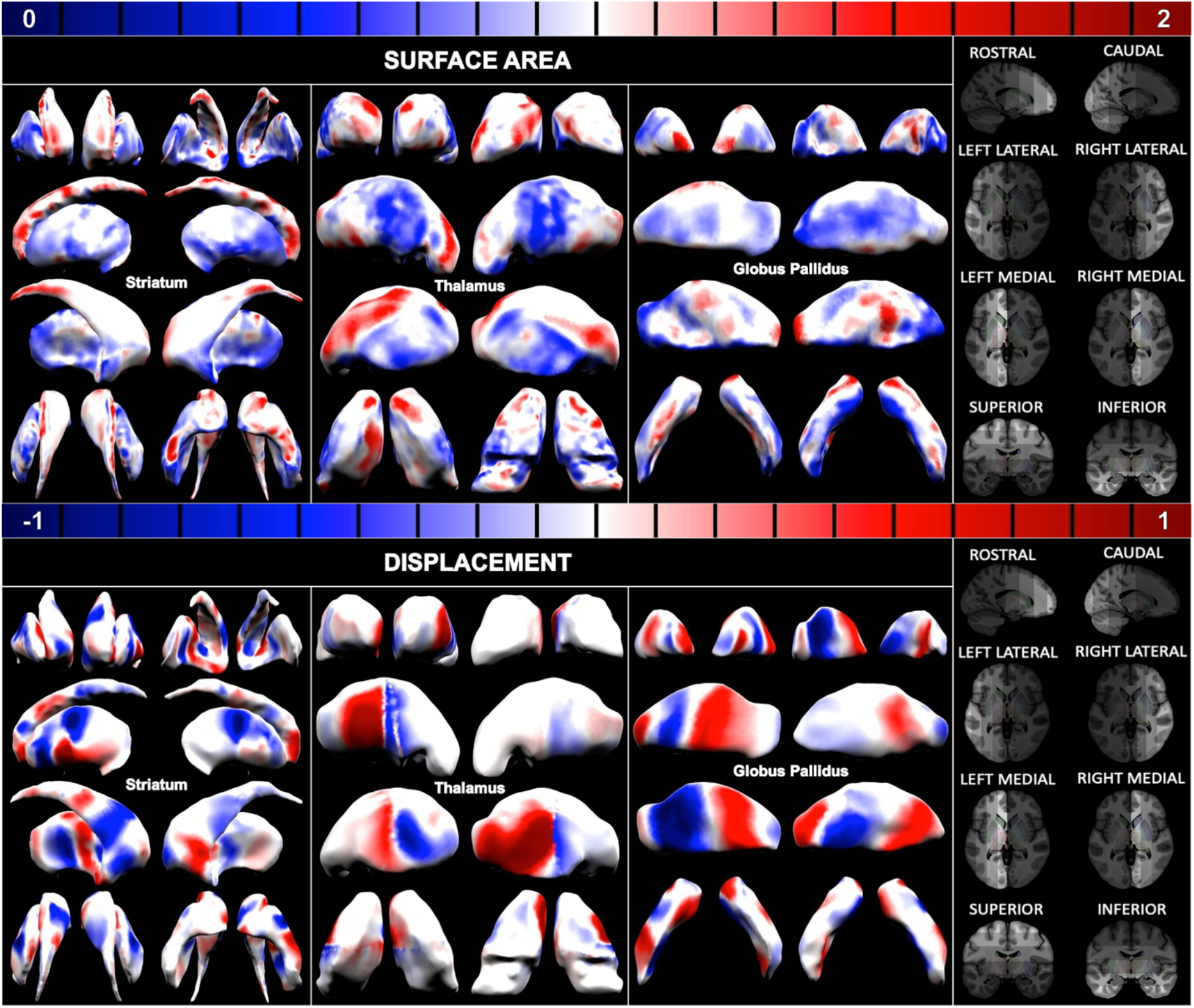
Interspecies differences in shape measures in the striatum, thalamus and globus pallidus. The top row shows the ratio of vertex-wise surface area measures in the human template to the same measures in the chimpanzee template. The bottom row shows the outward displacement going from the chimpanzee template to the human template. The views of the structures on display are shown in the column on the right-hand side; brighter areas on the brain indicate the view from which the structure is being seen.

When examining interspecies differences in the striatum between humans and chimpanzees, a very clear distinction between the caudate and putamen emerges: whereas the caudate area is expanded in humans, the putamen area is more conserved across species. We would therefore expect the caudate to have a higher human-to-chimpanzee volumetric ratio than the putamen. This was confirmed volumetrically, where we found that whilst the overall bilateral striatal volume ratio in humans to chimpanzees is of 1.97, it was of 2.34 for the caudate and 1.76 for the putamen (see Figure S2). We also found an anterior-posterior organization of alternatively conserved and expanded areas in the interspecies displacement differences between humans and chimpanzees.

Interspecies areal differences in the thalamus increase along a ventral-anterior to dorsal-posterior gradient. Additionally the area along the surface of the lateral thalamus is conserved in the center and expanded in small rostral and caudal areas. Interspecies displacement differences in the thalamus appear to be fairly asymmetrical: the caudal medial thalamus is expanded in the left hemisphere and conserved in the right hemisphere, whereas the opposite phenomenon occurs in the rostral medial thalamus. The caudal lateral thalamus is bilaterally expanded, but with more pronounced displacement differences in the left hemisphere.

The lateral globus pallidus is arealy conserved across species. It is outwardly displaced from chimpanzees to humans in a left anterior strip and small caudal patch of the left lateral globus pallidus and in a posterior strip of the right anterior globus pallidus. The medial globus pallidus has small patches of areal expansion. The internal segment is located along the medial-posterior globus pallidus and seems to be conserved in the left hemisphere and outwardly displaced in the right hemisphere.

### 2.2 Intraspecies analyses with the Human Connectome Project (HCP) dataset

The MAGeT Brain segmentation algorithm was used to automatically estimate subcortical volumes (Chakravarty *et al*., 2013; Pipitone *et al*., 2014; Makowski *et al*., 2017) and surface-based representations that were used to quantify vertex-wise surface area and surface displacement (Raznahan *et al*., 2014; Chakravarty *et al*., 2015; Shaw *et al*., 2015) on the high-quality MRI data provided by the HCP. After quality control across raw and processed data (see Figure S1) the overall sample was reduced to 821, 865, and 950 subjects for the striatum, thalamus and globus pallidus, respectively. See table S3 for demographic information.

#### 2.2.1 Heritability

Heritability is the proportion of variance of a trait that is attributable to genetics, as opposed to (unique or shared) environmental effects. We first examined phenotypes known to be highly heritable using a twin and non twin sibling model to verify if neuroanatomical phenotypes replicated previous studies. We observed high heritability across the structures of interest. The heritability estimates of TBV (82%) and left and right subcortical structure volumes, adjusted for sex and age, are high, as reported in previous studies (Blokland *et al*., 2012). Estimates range from 81% (left and right thalami and right globus pallidus) to 94% (right striatum) (see Figure 4). To assess the influence of overall brain size, we observed that adjusting for TBV increased heritability for all structure volumes by 1 to 3% aside from the left globus pallidus, which resulted in a 3% decrease (see Figure S3), implying that TBV has a negligible influence on subcortical volume heritability estimates.

**Figure 4.**
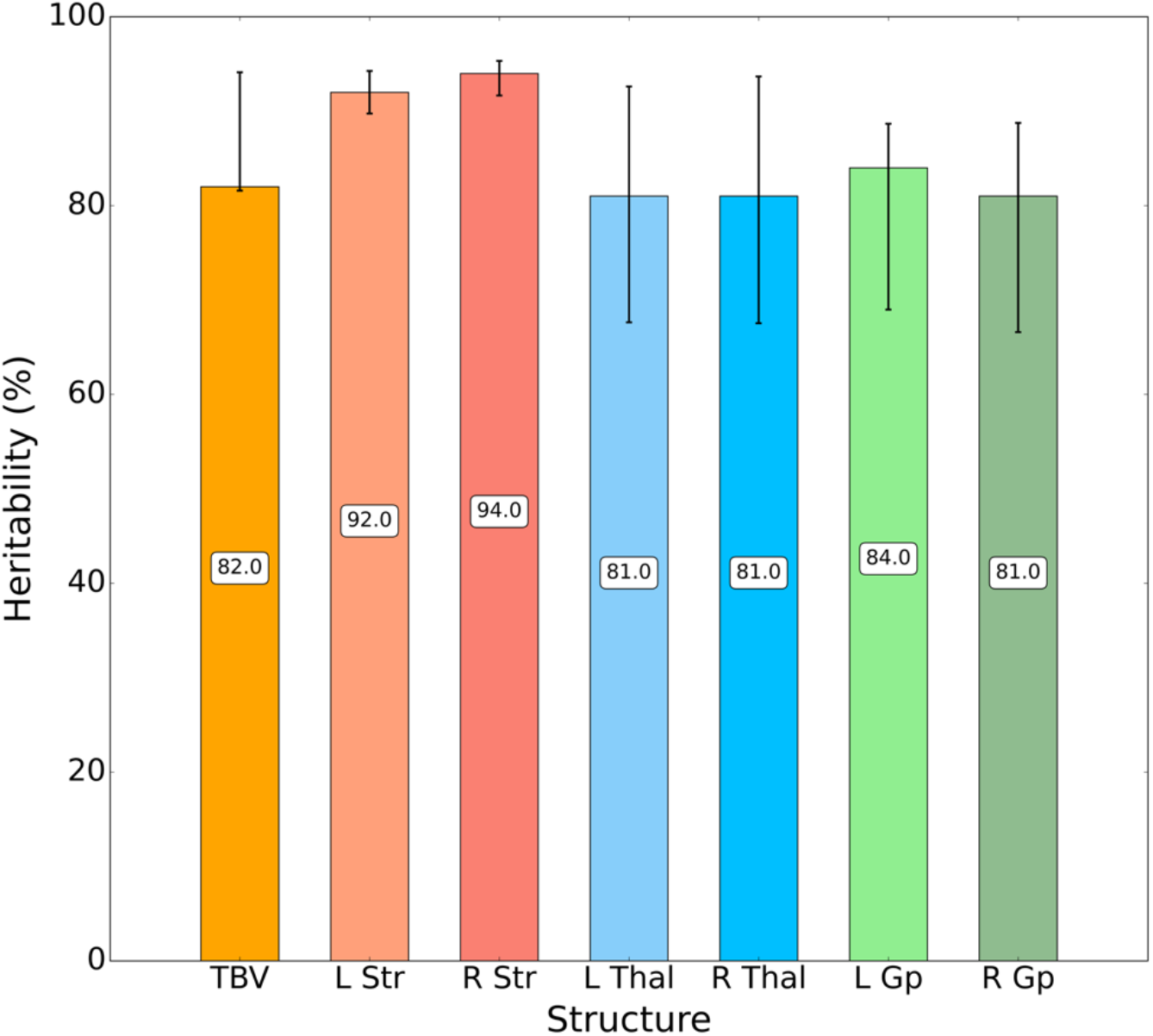
Heritability of TBV and ipsilateral structure-specific volumes, adjusted for sex and age. Shown for TBV and the left (L) and right (R) striatum (Str), thalamus (Thal) and globus pallidus (GP). Black bars indicate 95% confidence intervals.

We further examined heritability using a bivariate model to estimate the shared heritability (proportion of the covariance between two traits attributable to genetic effects) and genetic correlation (proportion of the genetic factors that control one trait which also control a second trait) between two phenotypes. We found that the shared heritability estimates between each of the left and right subcortical structure volumes and TBV are very high (ranging from 95% to 96%, see Figure S4). The genetic correlations between subcortical structure volumes and TBV range from 0.75 (right globus pallidus) to 0.97 (left thalamus) (see Figure S5).

To achieve a fine-grained topologically-specific surface-based mapping of regional heritability, we used the methods developed in recent studies demonstrating local surface area (or local measures of relative areal expansion or contraction (Raznahan *et al*., 2014; Shaw *et al*., 2015)) and displacement (local measure of bulging or indentation normal to the surface (Chakravarty *et al*., 2015; Voineskos *et al*., 2015)) as being relevant in the context of typical and atypical neurodevelopment. We adapted recent work from our group that maps vertex-level heritability of cortical thickness (Patel *et al*., 2018) to these measures (see Figure 5). The structure-specific bilateral average heritability estimates of sex and age-adjusted vertex-wise surface area and surface-based displacement measures are above 50%, suggesting that the variability of these traits is more strongly explained by genetic effects than shared or unique environmental effects (surface area mean heritability estimates: 68%, 66% and 63%; displacement mean heritability estimates: 62%, 57% and 65%; for the striatum, thalamus and globus pallidus, respectively). Heritability estimates increase along a lateral-medial gradient in the putamen and a dorsal-ventral gradient in the caudate for both surface area and displacement measures. Areal heritability estimates also increase along a posterior to anterior gradient along the thalamus. Vertex-wise displacement heritability is low in caudate and high in the putamen and estimates increase along an anterior and medial to postero-lateral gradient in the thalamus and a lateral-posterior to medial-anterior gradient along the globus pallidus.

**Figure 5.**
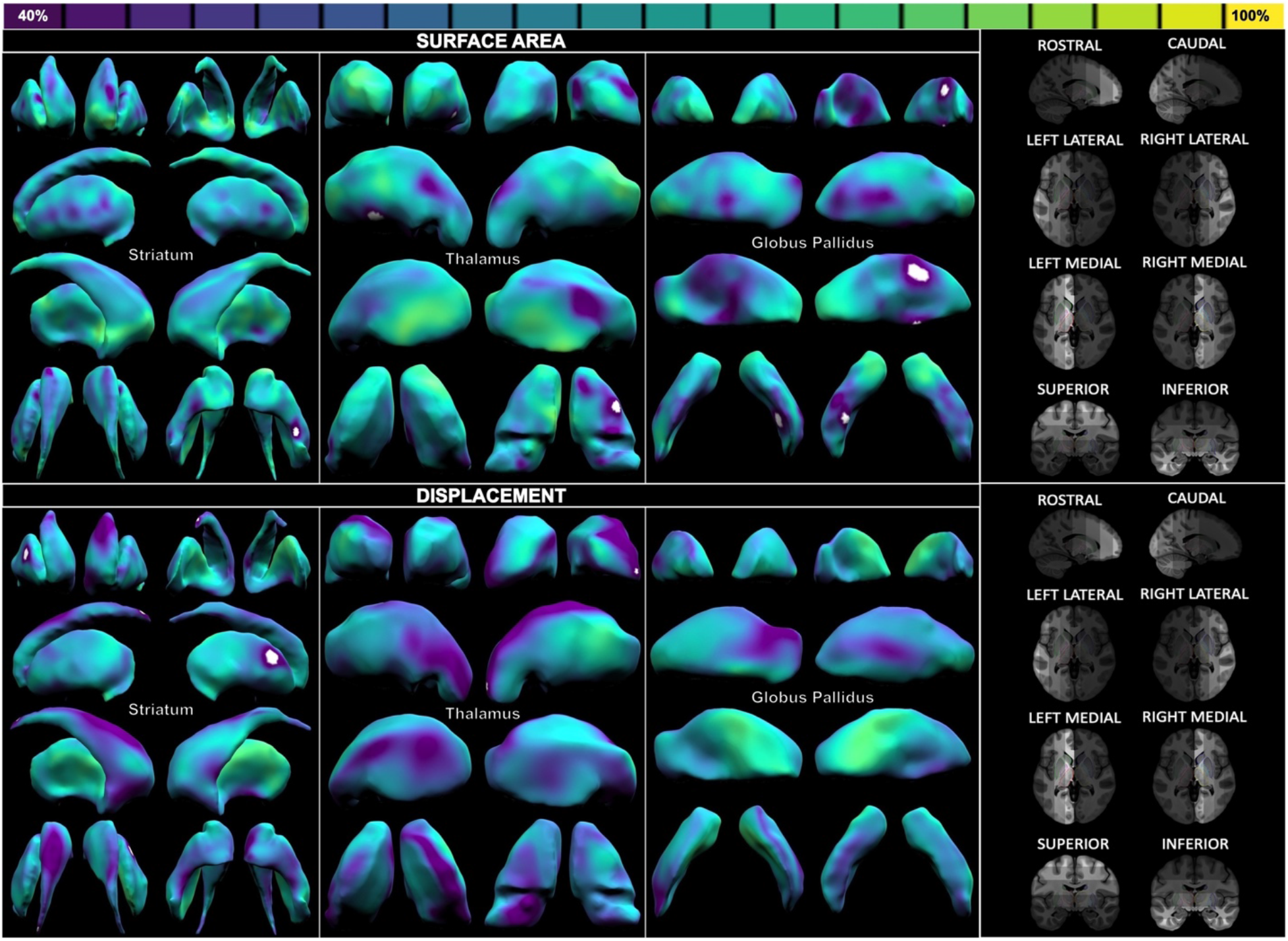
Heritability of sex and age-adjusted vertex-wise surface area (top row) and displacement (bottom row) in the striatum, thalamus and globus pallidus. The views of the structures on display are shown in the column on the right-hand side; brighter areas on the brain indicate the view from which the structure is being seen. Values in white are not significant (5% FDR correction). Ranges go from 40% (dark purple) to 100% (yellow).

We next adjusted vertex-wise measures for TBV before estimating heritability to determine whether the patterns we find are primarily driven by overall brain size effects (see Figure S6). Additional adjustment of vertex measures for TBV reduces areal heritability estimates by an average of 5% and does not affect the average heritability estimates of displacement measures. Qualitatively, heritability maps of TBV-adjusted vertex measures follow similar patterns to the ones observed in unadjusted measures. We adapted the cortical spin test (Alexander-Bloch *et al*., 2018) to the subcortical structures under examination to quantitatively compare the heritability maps of vertex-wise measures adjusted for TBV to those unadjusted for TBV and found the two sets of brain maps to be significantly and very strongly correlated (R=0.91, 0.89, 0.96 for surface area and R = 0.99, 0.99 and 0.99 for displacement, in the striatum, thalamus and globus pallidus), suggesting that our work captures highly spatially specific signatures of heritability that are not primarily driven by TBV effects.

#### 2.2.2 Partial least squares correlation (PLSC) analysis

We ran a PLSC analysis to relate subcortical vertex-wise measures extracted from the HCP dataset (brain variables) to a set of 30 behavioral attributes (associated with emotion, cognition and motor skill) and classified under different Instrument categories (see Figure 1.3). After performing a singular value decomposition of the covariance matrix between the two sets of variables (brain and behaviors), a PLSC analysis outputs significant latent variables (LVs) that relate patterns of neuroanatomy to patterns of behavior. We ran separate PLSC analyses for vertex-wise surface area and displacement.

Among the 2 significant LVs between vertex-wise surface area measures of the striatum, thalamus and globus pallidus and behavioral variables, the latent variable explaining most of the covariance between the two sets of variables loads predominantly onto higher-order cognitive behaviors (areal LV 1, 64% covariance explained) and another loads predominantly onto lower-order affective behaviors (areal LV 2, 7% covariance explained) (see Figure 6). More specifically, areal LV 1 is mainly associated with 3 measures of fluid intelligence and 2 measures of language whereas areal LV 2 is primarily associated with 3 measures of negative affect. The bootstrap ratio (BSR) of areal LV 1 is partially aliasing TBV (R = 0.59 between TBV and subject-specific brain scores; R = 0.21 between TBV and subject-specific behavior scores).

**Figure 6.**
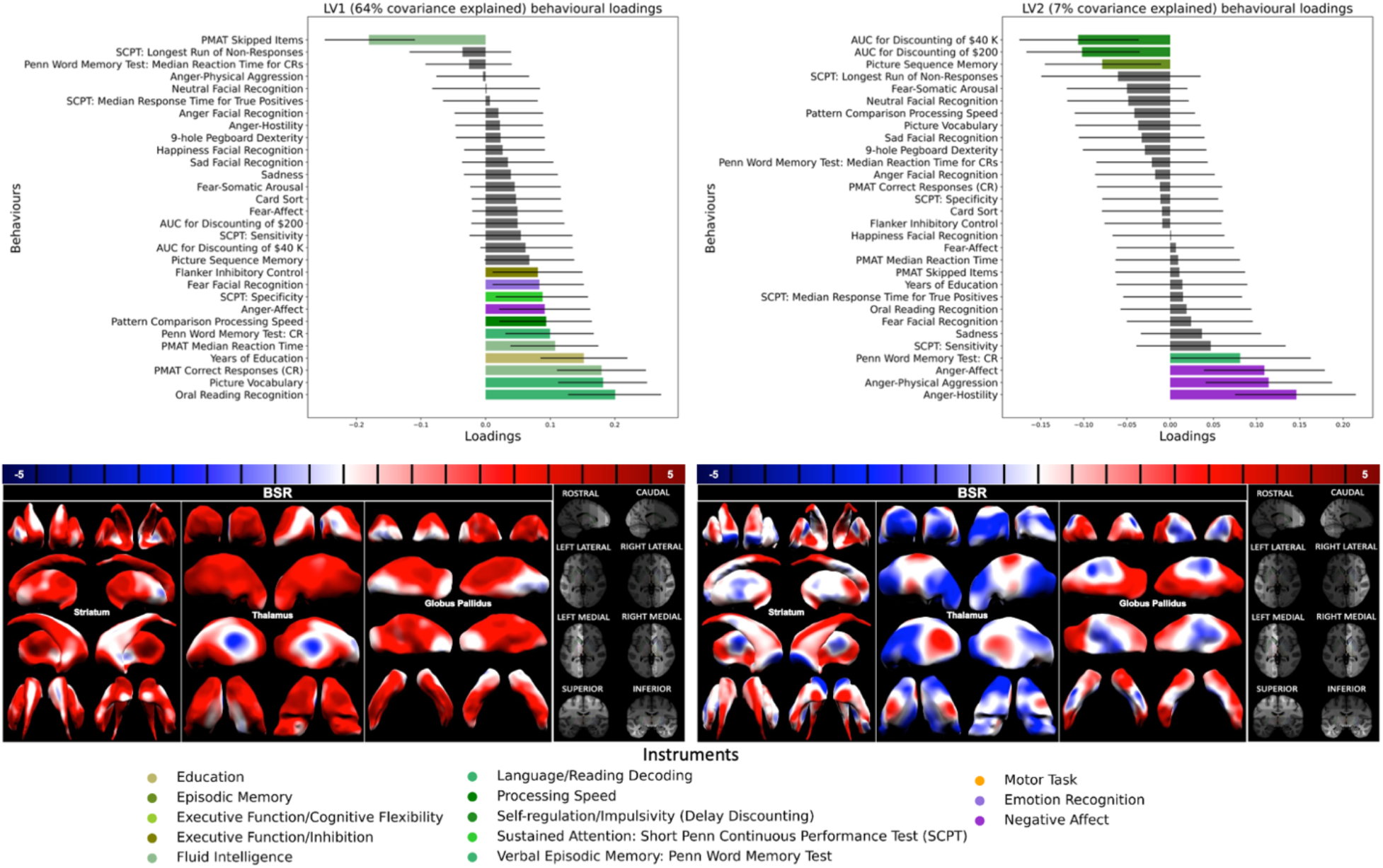
Areal LV 1 (left) and LV 2 (right). For each LV, the top row shows the behavior loadings and the bottom row shows the BSR. Behavior loadings whose 95% confidence intervals (white horizontal bars) have opposite signs are shaded in gray. Behavior measures are labeled on the y-axis and ordered such that loading value decreases from top to bottom. Behavior measures are classified by the HCP under different instruments and are therefore coloured by instrument in this figure (legend on the bottom), with shades of green, purple and orange corresponding respectively to cognition, emotion and motor instruments. BSR values (center row) are colored in red, white and blue and correspond respectively to values above 0, of 0 and below 0. The views of the structures on display are shown in the column on the right-hand side; brighter areas on the brain indicate the view from which the structure is being seen.

We additionally identified 4 significant LVs between vertex-wise displacement measures of the subcortex and behavioral variables (see Figure 7). Displacement LV 1 (26% covariance explained) is both cognitive and affective and loads onto 3 measures of fluid intelligence, 3 measures of facial recognition and 2 measures of language. Displacement LV 2 (12% covariance explained) and LV 3 (11% covariance explained) are both primarily affective: displacement LV 2 predominantly loads onto 2 measures of facial recognition and 2 measures of negative affect and displacement LV 3 predominantly loads onto 1 measure of facial recognition and 1 measure of negative affect. Finally, we found one primarily cognitive LV (displacement LV 4; 8% covariance explained) which loads onto the following instruments: cognitive flexibility, verbal episodic memory, inhibitory control, facial recognition and sustained attention.

**Figure 7.**
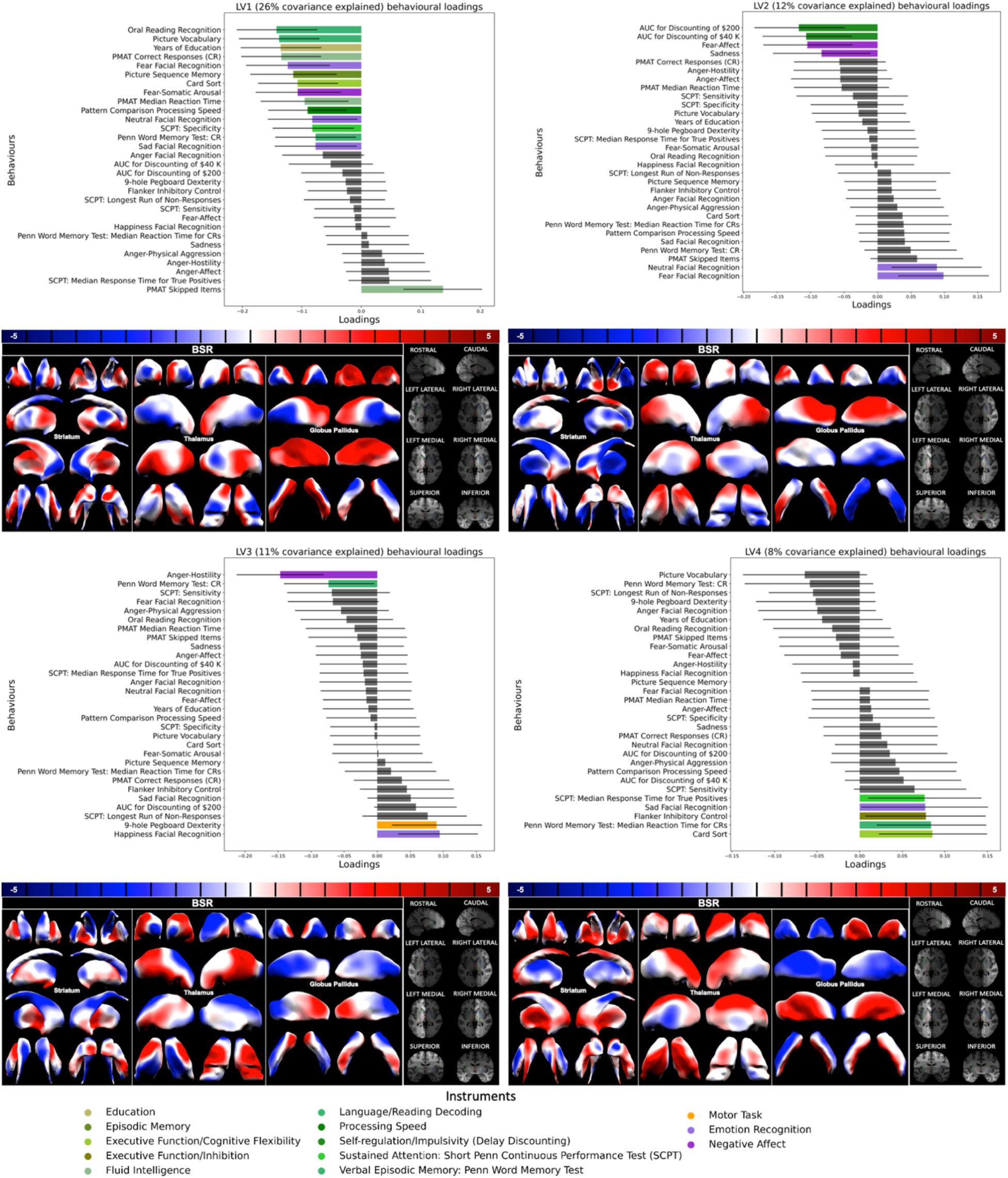
Displacement LV 1 (top left), LV 2 (top right), LV 3 (bottom left) and LV 4 (bottom right). Same legend as in Figure 6.

### 2.3 Cross-analysis brain map correlations

We were interested in examining the relationship between interspecies differences, heritability and latent variables of behavior in the striatum, thalamus and globus pallidus. We therefore adapted the surface-based cortical spin test (Alexander-Bloch *et al*., 2018) to the subcortical surfaces of interest in order to assess the significance of the spatial overlap (assessed via Pearson correlations) between pairwise combinations of the brain maps output by our first 3 analyses (interspecies difference maps, heritability maps and BSR maps) (see Figure 1.4). After applying 5% FDR correction across 42 comparisons, we found 33 significant correlations (see Figure 8).

**Figure 8.**
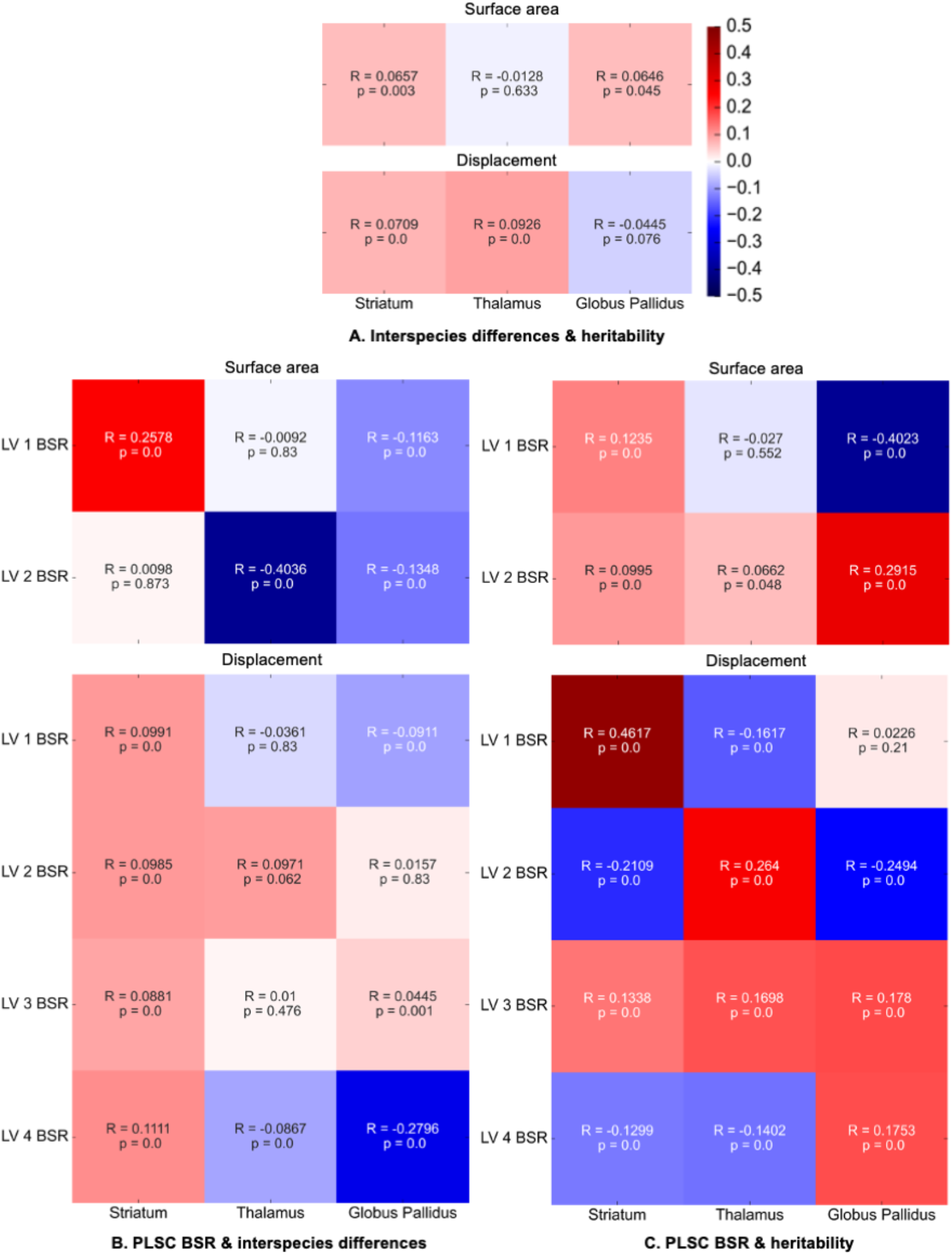
Pearson correlations and correlation significance between cross-analysis structure-specific brain maps. Panels A to C display the Pearson correlations between the following pairs of brain maps: heritability and interspecies difference maps (A), PLSC BSR and interspecies difference maps (B), and PLSC BSR and heritability maps (C). This figure also displays correlation significance (assessed via spin test) after FDR correction across 42 comparisons. The color map ranges from -0.5 to 0.5 and the color bar is located to the right of panel A. Striatal, thalamic and pallidal cross-analysis brain map correlations displayed in respectively the left, center and right columns of each panel. Vertex-wise surface area and displacement cross-analysis brain map correlations displayed in respectively the top and bottom row(s) of each panel.

#### 2.3.1 More heritable areas tend to be human-expanded

When directly comparing our heritability maps to our interspecies difference maps, we observed that more heritable areas are slightly more expanded in humans relative to chimpanzees. Indeed, there are 4 significant (p < 0.05) and modest positive correlations between heritability and interspecies difference maps in striatal area (R = 0.066), pallidal area (R = 0.065), striatal displacement (R = 0.071) and thalamic displacement (R = 0.093) (see Figure 8.A). However, areal LV 2 (cognitive and affective loadings) indirectly recapitulates what was previously observed in in the cortex (Patel *et al*., 2018; Reardon *et al*., 2018), as its BSR is positively correlated with heritability in the striatum (R = 0.10), thalamus (R = 0.066) and globus pallidus (R = 0.29) and negatively correlated with interspecies differences in the thalamus (R = -0.40) and globus pallidus (R = -0.13) (non-significant for the striatum).

#### 2.3.2 Human-expanded areas are associated with higher-order behaviors in the striatum

We observed interspecies differences in the striatum are positively correlated with cognitive areal LV 1 (R = 0.26) and cognitive displacement LV 4 (R = 0.11) (see Figure 8.B). Additionally, areal LV 1 and displacement LV 1, which primarily load onto language measures, are positively correlated with heritability in the striatum (R = 0.12 and R = 0.46, respectively) (see Figure 8.C).

#### 2.3.3 Areas that are conserved across species are associated with lower-order behaviors in the thalamus and globus pallidus

Interspecies differences are negatively correlated with the BSR of areal LV 2 and displacement LV 1 in the thalamus (R = -0.40 and R = -0.04, respectively) and globus pallidus (R = -0.13 and R = -0.10, respectively) (see Figure 8.B). Both areal LV 2 and displacement LV 1 load onto variables indexing affective behaviors (see Figure 8.C).

## 3. Discussion

Previous work examining the intersection of genetics (Patel *et al*., 2018), evolution and behavior (Reardon *et al*., 2018; Valk *et al*., 2020) has typically focused on the cerebral cortex. Here, we adapt many of the techniques previously used in this cortico-centric context to examine similar questions in the subcortical structures. A key innovation is the adaptation of the surface-based spin-test (Alexander-Bloch *et al*., 2018) to the striatum, thalamus and, globus pallidus to allow us to relate surface-based maps of chimpanzee-human differences, genetic influences, and complex behavioral phenotypes, and their inter-relationships.

Unlike the cortex (Patel *et al*., 2018; Reardon *et al*., 2018), we found that the subcortical regions that have the highest heritability are those that display the greatest areal expansion in humans relative to chimpanzees. The cortex and subcortex have been shown to follow different neurodevelopmental trajectories (Shaw *et al*., 2008; Hill *et al*., 2010; Raznahan *et al*., 2014). Whereas regions of cortical expansion that undergo the most pronounced growth in postnatal development overlap with those that differ the most between species in evolution (Krubitzer and Kahn, 2003; Hill *et al*., 2010), our results indicate a different pattern in the subcortical structures. These findings further stress the differing underlying mechanisms by which the subcortex is shaped throughout development and lifespan.

Additionally, the relationships between interspecies differences and behavior in the subcortex confirm our expectation that human-expanded areas would be associated with higher-order behaviors, whereas evolutionarily conserved areas would be associated with lower-order behaviors. In the striatum, we find significant positive correlations between 3 LVs that encode a set of higher-order cognitive variables and interspecies vertex-wise differences in the striatum. We also find 2 LVs loading predominantly onto affective variables to be negatively correlated with interspecies differences in the thalamus and globus pallidus. Finally, we observe 2 LVs that predominantly load onto human-specific language variables (often considered to be heritable (Canfield, Edelson and Saudino, 2017)) to be positively correlated with interspecies areal expansion and heritability maps, especially along the posterior putamen, a critical region for language function (Oberhuber *et al*., 2013; Viñas-Guasch and Wu, 2017).

Our results suggest that genetic processes mediating subcortical structures are crucial for higher-order behaviors in a way that is not reflected in the cortex. This further suggests that subcortical plasticity is shaped by analogous factors across species. Additionally, this work recapitulates the expectation that human-specific behaviors would be associated with more evolutionarily expanded areas in the subcortex, as can be observed in the cortex (Reardon *et al*., 2018).

### 3.1 Interspecies differences

In previous broad interspecies comparative studies, it has been shown that the allometric curve of neocortical volume relative to TBV is steeper than the allometric curve of striatal volume to TBV in primates (Striedter, 2006). In fact, recent work has shown that variation in primate brain structure scaling might be linked to socio-ecological adaptations (DeCasien and Higham, 2019). Consistent with these previous findings we found that while the ratio of human-to-chimpanzee TBV was 3.71, the ratios of subcortical volumes were lower, ranging from 1.52 (right globus pallidus) to 2.13 (left thalamus). Subcortical structures therefore contribute disproportionately less to TBV in humans relative to chimpanzees, as expected from the neurodevelopmental timing of different brain structures (Hill *et al*., 2010; Yopak *et al*., 2010; Raznahan *et al*., 2014). Areal expansion measures are likely most sensitive to relevant changes across species, as these differences maps are bilaterally symmetrical and not subject to residual affine effects.

Furthermore, we found an interesting trend in the striatum: the caudate is expanded in humans, whereas the putamen is conserved across species. The caudate plays an important functional role in two main cortico-subcortical circuits that begin and terminate in the prefrontal cortex (Alexander, DeLong and Strick, 1986), a structure that has been shown to be functionally reorganized in humans compared to other primates and displays a number of cellular and molecular evolutionary specializations (Eblen and Graybiel, 1995; Choi *et al*., 2017; Raghanti *et al*., 2018). The putamen is traditionally associated with limb and trunk movement (Kish, Shannak and Hornykiewicz, 1988), which can be thought of as lower-order behaviors. However, the putamen has been shown to be involved in orthographic processing (Oberhuber *et al*., 2013) (Viñas-Guasch and Wu, 2017) (Ford *et al*., 2013) and motor control-dependent speech production (Lieberman, 2002), which are human-specific functions. In fact, amino changes in the human-specific *FOXP2* gene have been associated with changes in dendritic branching, dopaminergic signaling, and neuronal physiology in the striatum which might be relevant to the acquisition of language function in humans (Enard *et al*., 2009).

In the thalamus, we observe that the surfaces of structures involved in visual information processing, the limbic, and two prefrontal cortico-subcortical loops seem to be expanded in humans relative to chimpanzees, whereas the surfaces of structures involved in mechanoreception and motor control are more conserved between these species. For example, the anterior nucleus (Papez, 1937) and posterior-medial medial dorsal nucleus (Alexander, DeLong and Strick, 1986) of the limbic system are relatively expanded in humans compared to chimpanzees. The pulvinar nucleus, a component of the oculomotor circuit (Arend *et al*., 2008), and portions of the anterior nucleus, involved in visual information processing and two prefrontal loops (Alexander, DeLong and Strick, 1986), are also relatively expanded. On the other hand, the ventral posterolateral region, which relays information associated with mechanoreception (Armstrong, 1976), and the ventral lateral nucleus, involved in motor control, are conserved between humans and chimpanzees.

The pallidal external segment, which modules striatal inputs to the internal segment (Purves *et al*., 2018) is more areally conserved across species than the internal segment, which is directly involved in all of the cortico-subcortical circuits (Alexander, DeLong and Strick, 1986).

### 3.2 Heritability

Our heritability estimate for TBV falls within the 95% confidence intervals of previous studies (Blokland *et al*., 2012), (Gómez-Robles *et al*., 2015) (Kremen *et al*., 2010) (Patel *et al*., 2017) (Wen *et al*., 2016) and, consistent with previous literature, we found that TBV is more heritable than subcortical structure volume (Blokland *et al*., 2012). Heritability estimates of subcortical shape measures were, on average, lower than volumetric heritability estimates, a finding that has previously been shown in the caudate (Ge *et al*., 2016), putamen (Roalf *et al*., 2015), nucleus accumbens (Roalf *et al*., 2015) and thalamus (Ge *et al*., 2016) (Roalf *et al*., 2015). Our results are consistent with previous findings demonstrating that adjusting region-specific measures for aggregate measures yields slightly lower heritability estimates. Indeed, work in the cortex has found that adjusting vertex-wise cortical thickness for average ipsilateral cortical thickness and vertex-wise cortical surface area for TBV results in lower heritability estimates across the cortical mantle (Patel *et al*., 2018). Similarly, we found that adjusting vertex-wise measures for TBV lowers heritability estimates for both surface area and displacement. Furthermore, as has been observed of cortical thickness and surface area (Patel *et al*., 2018), heritability estimates are slightly higher for vertex-wise areal measures than vertex-wise displacement measures. However, in our work, adjusting vertex-wise measures for TBV did not affect heritability estimates by more than 5% and did not impact heritability patterning along our subcortical surfaces, suggesting that our results capture highly spatially specific signatures of subcortical heritability.

Interestingly, we observe high heritability estimates in neuroanatomical regions that have been previously associated with highly heritable disorders or behaviors. For example, Huntington’s Disease, a genetically determined disorder (Ross *et al*., 2014), is marked by cellular loss in the caudate head (Vonsattel *et al*., 1985), a highly heritable region. Additionally, elevated heritability estimates of displacement measures along the surfaces of structures involved in the motor loop (Alexander, DeLong and Strick, 1986), namely the putamen, ventral lateral thalamic nucleus and ventro-lateral pallidal internal segment, converge with the behavioral literature, as motor learning and control have been shown to be heritable (Missitzi *et al*., 2013). High areal heritability estimates along the surface of the nucleus accumbens, involved in emotion recognition (Satterthwaite *et al*., 2011), are consistent with observations in addiction (Di Chiara *et al*., 2004; Li and Burmeister, 2009), another disorder under strong genetic control (Scofield *et al*., 2016). We cautiously interpret these findings as a suggestion that topological specificity may provide relevant information regarding neuropsychiatric pathophysiology.

Finally, we observed a trend in both areal and displacement measures where the lateral pallidal internal segment is generally less heritable than the external segment. Given that the internal segment integrates projections from both the external segment and from the striatum (Purves *et al*., 2018), it is cumulatively impacted by a wider array of inputs than the external segment, which could in turn make the internal segment more plastic and sensitive to the influence of non-genetic environmental factors.

### 3.3 Brain-behavior dimensions

In our multivariate analyses examining brain and behavior we demonstrate sensitivity to the anatomico-behavioral covariance through the series of LVs that emerged. We found that the anatomical patterning of a latent variable related to high-order cognition (areal LV1) increases along an anterior to posterior gradient along the putamen. Behaviorally, this LV loads on variables related to language performance (oral reading recognition and picture vocabulary task) and is consistent with previous work demonstrating the differential sensitivity to actual words in the anterior versus posterior putamen (Oberhuber *et al*., 2013). We found a second significant LV (areal LV 2) that is mostly associated with measures of negative affect and delay discounting and which loads strongly onto the medial-rostro-dorsal caudate, a region where cortical inputs associated with reward, cognitive control and attention converge (Choi *et al*., 2017).

For our PLSC analysis where we input vertex-wise displacement measures as our set of brain variables, we found one cognitive and affective LV (displacement LV 1), one primarily cognitive LV (displacement LV 4) and two primarily affective LVs (displacement LVs 2 and 3). Displacement LV 2, which loads onto many affective variables, has mostly negative loadings along the entire striatum aside from the ventral striatum, where the loadings are strongly positive. This is interesting given the involvement of the nucleus accumbens in the subcortical-cortical limbic loop (Alexander, DeLong and Strick, 1986) and in emotion regulation (Satterthwaite *et al*., 2011). On the other hand, the primarily cognitive LV has strong positive loadings along the medial rostral-dorsal caudate, which integrates higher-order cognitive inputs (Choi *et al*., 2017).

### 3.4 Cross-analysis brain map correlations

When directly examining the relationship between heritability and interspecies differences maps, we found that more heritable areas are slightly more expanded across species, a finding that runs counter to what has been observed in the human cerebral cortex (when compared to macaques) (Patel *et al*., 2018) (Reardon *et al*., 2018). Nevertheless, we observe an interesting trend and our correlations withstand a surface-based spin test (Alexander-Bloch *et al*., 2018) and multiple comparison correction (Benjamini and Hochberg, 1995).

Additionally, by comparing BSR maps and interspecies difference maps we observe that evolutionarily expanded portions of the striatum are associated with cognitive measures whereas evolutionarily conserved regions of the thalamus and globus pallidus are associated with affective measures, in line with our expectations regarding the relationship between behavior and evolutionary neuroanatomical divergences. More specifically, the cognitive LVs that positively correlate with interspecies differences in the striatum are also positively correlated with heritability estimates. These LVs load onto surface measures of the putamen and language-related behaviors which are a human-specific, heritable (Stromswold, 2001) and previously associated with brain activity in the putamen (Oberhuber *et al*., 2013; Viñas-Guasch and Wu, 2017). On the other hand, in the thalamus and globus pallidus, we found that the BSR values of 2 affective behaviors are higher in regions that are more conserved between humans and chimpanzees. Areal LV 2 is mainly driven by affective measures pertaining to anger, which has previously been associated with thalamic activity (Kimbrell *et al*., 1999; Damasio *et al*., 2000; Fabiansson *et al*., 2012).

Finally, we cautiously speculate that some correlation directionalities may be associated with connectivity to other brain structures. There are sets of correlations (BSR of areal LV 1 and heritability; BSR of displacement LV 4 and heritability) that go in an opposite direction in the globus pallidus (not directly connected to the cortex) relative to the striatum and thalamus (directly connected to the cortex). Furthermore, in the primarily affective displacement LV 2, correlation magnitudes are all higher than 0.2 and correlation directionality can tentatively be attributed to connectivity with the amygdala, as the BSR of this LV is positively correlated with heritability in the thalamus, which shares direct connections with the amygdala, and negatively correlated with heritability in the striatum and globus pallidus, which do not connect with the amygdala.

### 3.5 Limitations

Firstly, heritability estimation, like any statistical model, relies on a simplification of reality that is subject to certain limitations. For instance, previous work has shown that gene-environment interactions lead to overestimation of heritability as they are often confounded for additive genetic effects (Dalmaijer, 2020). Secondly, parental passive genetic influence results in the marginal under-estimation of shared environmental effects and measurement error increases the estimation of non-shared environmental effects (Dalmaijer, 2020). Thirdly, although the purpose of our comparative neuroanatomy work was to try to better understand human brain evolution, it should be acknowledged that pairwise contrasts with only chimpanzees cannot fully take allometric scaling into account, nor can chimpanzee brain structure be assumed to perfectly represent the ancestral condition from which human neuroanatomy evolved (Staes *et al*., 2019). Furthermore, interspecies differences between humans and their closest phylogenetic relatives (Perelman *et al*., 2011) are still but a proxy for a research question that would require longitudinal data on an inaccessible time scale.

## 3.6 Conclusion

Previous literature has shown a strong convergence between functional and evolutionary expansion maps of the cortex (Ardesch *et al*., 2019; Wei *et al*., 2019; Sydnor *et al*., 2021). Additionally, evolutionarily expanded heteromodal cortical areas, associated with higher-order cognitive behaviors, have been shown to be less heritable than more conserved unimodal cortical areas, associated with lower-order cognitive behaviors (Patel *et al*., 2018). We examined these previously unknown relationships in the subcortex (striatum, thalamus and globus pallidus), using surface-based brain maps of heritability, interspecies differences and latent variables of behavior.

We found many significant correlations between pairs of brain maps. Importantly, we found that more heritable areas are more expanded across species and evolutionary expanded and conserved areas are respectively associated with cognitive and affective behavioral measures. Overall, the purpose of examining fine-grained surface-based shape measures is to better infer the underlying cellular distribution of these structures than volumetric measures could ever convey. Future work should aim to examine how these maps converge with subcortical cytoarchitecture and gene expression.

## Acknowledgments

M.M.C. receives salary support from the Fonds de recherche du Québec (FRQS) and research funding from the National Sciences and Engineering Research Council of Canada (NSERC), the Canadian Institutes of Health Research (CIHR) and Healthy Brains for Healthy Lives (HBHL). N.B. has received scholarships from HBHL and Canadian Graduate Scholarships – Master’s (CGS M) to support her studies.

## Author Contributions

M.M.C. conceptualized and directed the study. N.B. executed the project. G. A. D. generated the chimpanzee MRI template, conceptualized the interspecies MRI registration protocol and provided technical support. S. P. contributed the baseline heritability code. R. P. contributed to the scripting necessary for high-performance computing. S. P., R. P., S. T., E. P. and S. A. B showed N.B. laboratory protocols (analysis, image processing, quality control). M. C. and R. M. contributed to the PLSC code. O. P. contributed to the spin test code. C. C. S, W. D. H., J. S. and A. R. contributed the chimpanzee data and provided significant feedback.

## STAR Methods

### 1. Resource availability

#### Lead contact

Further information and requests for resources should be directed to and will be fulfilled by the lead contact, M. Mallar Chakravarty.

#### Materials availability

This study did not generate new unique reagents.

#### Data and code availability

- This paper analyzes existing, publicly available data. These accession numbers for the datasets are listed in the key resources table.
- All original code has been deposited at [repository] and is publicly available as of the date of publication. DOIs are listed in the key resources table.
- Any additional information required to reanalyze the data reported in this paper is available from the lead contact upon request

## 2. Experimental model and subject details

### 2.1 Interspecies analysis: human average MRI template

For the human magnetic resonance imaging (MRI) template, the openly available 0.3 mm resolution 3T T1w brain template from the Multiple Automatically Generated Templates brain segmentation algorithm (MAGeT) brain segmentation algorithm pipeline (https://github.com/CoBrALab/MAGeTbrain) was selected. This template was made by averaging the high-resolution 3T T1w MRI data from 5 subjects (3 females, 2 males, mean age 37 years, age range 29-57 years) (Winterburn *et al*., 2013; Park *et al*., 2014; Tullo *et al*., 2018).

### 2.2 Interspecies analysis: chimpanzee average MRI template

For the chimpanzee MRI template, a 0.4 mm isotropic T1w template created by averaging the chimpanzee MRI data from the National Chimpanzee Brain Resource (NCBR) using Advanced Normalization Tools (ANTS) (Avants, Tustison and Song, 2009) was selected (Devenyi *et al*., 2021). The MRI sample consisted of 139 1.5 T and 77 3T MR images. Of the 216 chimpanzees, 129 of them were female and the average age was 27.

### 2.3 Intraspecies analyses: Human Connectome Project (HCP) dataset

Structural MRI data was acquired from the WU-Minn Human Connectome Project (HCP) S1200 Release (Van Essen *et al*., 2013; Glasser *et al*., 2016). The HCP consortium is an open-data initiative that was launched in 2010 with the aim to propel the investigation of brain connectivity in a dataset that captures the ethnic, racial, behavioral and economic demographic variability of the United States (Van Essen *et al*., 2013). Brain imaging, behavioral and genotyping data on 1,200 healthy young adult twin and non-twin siblings, aged 22-35, was gradually released between 2013 and 2017.

Participants with a significant history of psychiatric disorder, substance abuse (hospitalization for the condition for two days or longer), neurological disease, cardiovascular disease or genetic disorder were excluded from the HCP. Premature birth (before 34 weeks for twins; before 37 weeks for non-twin siblings; less than 5 lbs at birth for non-twin siblings if weeks unknown) was also an exclusion criteria. Smokers, recreational drug users and participants who were overweight were included, in order to better capture population diversity. After participants were screened for exclusion criteria, participants left without a sibling were also excluded from the project. See Table S1 of (Van Essen *et al*., 2013) for more information on the criteria for inclusion and exclusion of the HCP participants.

In March 2017, imaging, behavioral and genotyping data on the final group of subjects was released. The S1200 Release (Reference Manual: https://www.humanconnectome.org/storage/app/media/documentation/s1200/HCP_S1200_Release_Reference_Manual.pdf) includes 1,206 subjects 1113 of which have structural magnetic resonance imaging (MRI) scans. Structural images were collected using a Siemens 3-Tesla (3T) Skyra scanner that was modified with a Siemens SC72 gradient coil from 40 mT/m to 100 mT/m in order to increase maximum gradient strength (Van Essen *et al*., 2012, 2013). The current study uses the high-resolution 3T T1-weighted (T1w) MRI images (TE/TR=2.14/2400 ms, TI=1000ms, a=8°, 0.7 mm^3^ isotropic voxels, HCP S1200 Reference Manual) from the S1200 subject release (n = 1086 subjects, HCP S1200 Reference Manual, downloaded directly from the HCP online repository (https://db.humanconnectome.org/data/projects/HCP_1200)).

We obtained consent to access the HCP restricted behavioral attributes (https://wiki.humanconnectome.org/display/PublicData/HCP-YA+Data+Dictionary-+Updated+for+the+1200+Subject+Release). Each attribute, measured in every participant, belongs to an instrument category. Many attributes correspond to cognitive tests developed at the National Institutes of Health (NIH) or at the University of Pennsylvania (Penn). We selected the following instruments and attributes (listed in parenthesis) for this study: demographics (sex, age, years of education), episodic memory (NIH Toolbox Picture Sequence Memory Test: Unadjusted Scale Score), executive function and cognitive flexibility (NIH Toolbox Dimensional Change Card Sort Test: Unadjusted Scale Score), executive function and inhibition (NIH Toolbox Flanker Inhibitory Control and Attention Test: Unadjusted Scale Score), fluid intelligence based on a subject’s correct responses out of a total of 24 questions (Penn Progressive Matrices: Number of Correct Responses, Total Skipped Items, Median Reaction Time for Correct Responses), language and reading decoding (NIH Toolbox Oral Reading Recognition Test: Unadjusted Scale Score), language and vocabulary comprehension (NIH Toolbox Picture Vocabulary Test: Unadjusted Scale Score), pattern completion processing speed (NIH Toolbox Pattern Comparison Processing Speed Test: Unadjusted Scale Score), self-regulation and impulsivity (Delay Discounting: Area Under the Curve for Discounting of $200, Area Under the Curve for Discounting of $40,000), sustained attention (Short Penn Continuous Performance Test: Median Response Time for True Positive Responses, Sensitivity, Specificity, Longest Run of Non-Responses), verbal episodic memory (Penn Word Memory Test: Total Number of Correct Responses, Median Reaction Time for Correct Responses), dexterity (NIH Toolbox 9-hole Pegboard Dexterity Test: Unadjusted Scale Score;), Emotion Recognition (Penn Emotion Recognition Test: Number of Correct Anger Identifications, Number of Correct Fear Identifications, Number of Correct Happy Identifications, Number of Correct Neutral Identifications, Number of Correct Sad Identifications), Negative Affect (NIH Toolbox Fear-Somatic Arousal Survey: Unadjusted Scale Score, NIH Toolbox Anger-Affect Survey: Unadjusted Scale Score, NIH Toolbox Anger-Hostility Survey: Unadjusted Scale Score, NIH Toolbox Anger-Physical Aggression Survey: Unadjusted Scale Score, NIH Toolbox Sadness Survey: Unadjusted Scale Score).

Manual quality control (see Figure S1) and removal of subjects without a sibling reduced the sample size to 950 subjects, including 517 females, 433 males, 260 monozygotic twins, 156 dizygotic twins and 534 non-twin siblings. The average age of the sample was 28.80 years ± 3.65 SD (range 22-35 years). See Tables S1 and S2 for more information on the subject demographic information and behavioral attributes included in this study.

## 3. Method Details

### 3.1 Warping subcortical surface files from the human MRI template to the chimpanzee MRI template

We used an adapted group-wise strategy (https://github.com/ANTsX/ANTs) (Avants, Tustison and Song, 2009) to create a mean subcortical human-chimpanzee hybrid template (see Figure 1.1.B) representing the minimum voxel-wise deformation that is required to align the subcortical structures across species. Registration estimation was constrained to the subcortical structures of interest (see Figure 1.1.A) as a means of improving registration accuracy (Chakravarty *et al*., 2008, 2009). The MAGeT subcortical surfaces (https://github.com/CobraLab/atlases/tree/master/5-atlas-subcortical) (Tullo *et al*., 2018) were transformed from the human to the chimpanzee, using the vector warp fields generated by ANTS when creating the human-chimpanzee hybrid (see Figure 1.1.C). Since ANTS registration outputs a set of linear and nonlinear transformations, we only used the non-linear transformations in order to account for alignment and volumetric differences between the human and chimpanzee template MRIs. Additionally, for the displacement measures, we used ANTS in order to remove any residual affine effects from our vector fields to make sure to be capturing displacement differences that are not driven by translation effects.

### 3.2 Human Connectome Project (HCP) data processing

#### 3.2.1 Image preprocessing

Structural MR images were acquired after minimal preprocessing by the HCP PreFreeSurfer pipeline. Briefly, the PreFreeSurfer pipeline involves gradient distortion correction, ACPC alignment and readout distortion correction of the T1w images (Glasser *et al*., 2013). After conversion from NIFTI to MINC format (see Figure 1.2/3.A), structural MR images were further preprocessed in our lab using the minc-bpipe library (https://github.com/CobraLab/minc-bpipe-library) default preprocessing pipeline: N4 correction for intensity non-uniformity (Tustison *et al*., 2010), cropping of the neck region and brain mask generation using brain extraction based on nonlocal segmentation technique (BEaST) (Eskildsen *et al*., 2012) (brain masks were used for total brain volume calculation) (see Figure 1.2/3.B). Manual quality control for minc-bpipe outputs and motion reduced the sample size to 1033 subjects (see Figure S1). Removal of subjects without siblings reduced the sample size to 950 subjects.

#### 3.2.2 Image preprocessing: manual quality control ratings

Subjects were rated for motion on a scale of 1 to 4 (https://github.com/CoBrALab/documentation/wiki/Motion-Quality-Control-(QC)-Manual) (Bedford *et al*., 2019): a score of 1 was attributed to images where the motion artifacts were negligible; a score of 2 was attributed to images where there was slight but visible ringing (mostly near the cortical surface); a score of 3 was attributed to images where at least a third of 2-3 slices had significant ringing; a score of 4 was attributed to images where at least a third of the slices had significant ringing. Subjects with a score of 2 or higher were excluded from the study. None of the subjects from the study sample had a score higher than 3. See Figure S1 for rating examples.

#### 3.2.3 Image segmentation

Volumetric and surface-based morphometry measures for subcortical structures were derived using the publicly available MAGeTbrain algorithm was used (https://github.com/CobraLab/MAGeTbrain) (Chakravarty *et al*., 2013; Pipitone *et al*., 2014; Makowski *et al*., 2018) using subcortical definitions defined using an atlas based on serial histological sections (Chakravarty *et al*., 2006) and warped to fit 5 high-resolution MRI-atlases (Tullo *et al*., 2018) (https://github.com/CobraLab/atlases/tree/master/5-atlas-subcortical) (see Figure 1.2/3.C).

First, 21 randomly selected template brains were labeled using the 5 high-resolution MRI-atlas and all six subcortical labels at once (bilateral striatum, thalamus, and pallidum). The template brains were propagated to each subject brain (including the original templates), yielding 105 segmentations per subject. For each subject, these segmentations were fused via voxel-by-voxel majority voting. This multi-atlas-based segmentation approach has been shown to better account for the neuroanatomical diversity of the sample under study (Chakravarty *et al*., 2013; Tullo *et al*., 2018). The accuracy of this methodology has been previously validated across several studies (Chakravarty *et al*., 2013; Makowski *et al*., 2018; Tullo *et al*., 2018).

We improved the final segmentation accuracy (based on multiple runs of manual quality control by N.B.) by taking a region-of-interest-based approach for all nonlinear registrations performed. We have previously demonstrated that this approach improves segmentation accuracy (Chakravarty *et al*., 2008, 2009). For each structure we used the 21 “best” templates as per our previous observations. To achieve improved segmentations, N.B. corrected a thalamic under-segmentation originally observed in the original 5 input atlases being used (Tullo *et al*., 2018). The final sample size after quality control were 821, 865, and 950 subjects for the striatum, thalamus, and pallidum, respectively.

For all of the HCP subjects, the human MRI template and the chimpanzee MRI template, total brain volume and ipsilateral volumes for the striatum, thalamus and globus pallidus were obtained from the brainmask (total brain volume) or structure-specific labels.

#### 3.2.4 Image segmentation: manual quality control ratings

Segmentations were rated from 0 to 1 (https://github.com/CoBrALab/documentation/wiki/MAGeT-Brain-Quality-Control-(QC)-Guide): a score of 1 indicated that the image segmentation had negligible errors (less than 5 slices with minor errors), a score of 0.5 indicated that the image segmentation had errors but still passed (minor errors in 5-10 slices) and a score of 0 indicated that the image had to be excluded from the study (major errors in 2-3 slices or minor errors in over 10 slices). See Figure S1 for examples of over–, under– and correct segmentation across the structures of interest.

#### 3.2.5 Obtaining vertex-wise shape measures

We estimated vertex-wise shape measures to estimate morphological variation as in our previous work (Raznahan *et al*., 2014; Shaw *et al*., 2014, 2015; Chakravarty *et al*., 2015; Makowski *et al*., 2018) (see Figure 1.2/3.D). The final outputs provide a one-to-one vertex correspondence across subjects that allow for group-wise statistical analyses of shape measures. Previously, our group has generated surface meshes for the striatum (13 000 vertices), thalamus (6500 vertices) and globus pallidus (3000 vertices) on the MAGeT brain template brain (Tullo *et al*., 2018), that we are using here to represent a canonical human anatomical space. These surface meshes can be propagated from the MAGeT brain template brain through the chain of transformations to each individual subject. The 105 resultant surfaces are then fused by positioning the vertex-wise median coordinate position (Raznahan *et al*., 2014; Chakravarty *et al*., 2015; Shaw *et al*., 2015). To compute vertex-wise surface area, a Voronoi parcellation using 3 seeds per triangle face assigns 1 seed to each vertex (Boots *et al*., 2009)(Raznahan *et al*., 2014). To compute vertex-wise displacement on one subject, 105 deformation fields (vectors going from model, to atlas, to template, to subject) are averaged to create a deformation vector. The displacement is then computed as the projection of the final deformation vector at each vertex onto the unit vector of the surface normal to the original surface at that vertex (Lerch *et al*., 2008; Voineskos *et al*., 2015). The surface-based measures are blurred with a surface-based diffusion-smoothing kernel of 5 mm for the striatum and thalamus, and 3 mm for the globus pallidus (Kim *et al*., 2005).

## 4. Quantification and statistical analysis

### 4.1 Interspecies analysis: vertex-wise shape differences between humans and chimpanzees

Vertex-wise surface area maps were extracted with the depth_potential function (within the CIVET 2.0 package (Ad-Dab’bagh *et al*., 2006), and the expansion map between two brains was created by examining the ratio between template-specific vertex-wise surface area, after accounting for linear differences (see STAR Methods 3.2) (see Figure 1.1.D). Vertex-wise displacement was obtained by computing the dot product of the vertex surface normals and the deformation vector field, yielding estimates of the direction and magnitude of the inward or outward displacement of the surface, using the object_volume_dot_product code (https://github.com/Mouse-Imaging-Centre/minc-stuffs/blob/master/src/object_volume_dot_product.c). Interspecies differences between the displacement values were examined after accounting for linear differences (see STAR Methods 3.2) (see Figure 1.1.D).

### 4.2 Intraspecies analysis: heritability estimation in the Human Connectome Project (HCP) dataset

We used a similar strategy for heritability estimation as our previous work on hippocampal subfield volumes (Patel *et al*., 2017) and cortical morphometry (Patel *et al*., 2018) (see Figure 1.2.E). Heritability was estimated using a twin and non-twin sibling design and the OpenMx package (version 2.12.2) (Boker *et al*., 2011) in R (version 3.5.1). A structural equation model (SEM) partitioned the phenotypic variance (**σ**^2^) of the trait under study into three components: genetic effects (A), common environment (C) and unique environment (E) (such that **σ**^2^ = **σ**_A_^2^ +**σ**_C_^2^+**σ**_E_^2^). Broad-sense heritability (H^2^) is defined as the proportion of variance of the studied phenotype that is attributable to genetic effects, meaning the ratio between the genetic effects (A) and the total variance (**σ**^2^) (Mayhew and Meyre, 2017).

The A is set to 1 between siblings in monozygotic twin pairs and 0.5 in dizygotic twin pairs or non-twin siblings, reflecting the assumption that 100% and 50% of the genotype is shared in each group, respectively. Common environment is assumed to be the same for all siblings, while unique environment (E) does not correlate between siblings. The significance of the estimator for variance attributable to genetic effects (**σ**_A_) is assessed by comparing the original ACE model to an AE model (a model where the genetic effects parameter was removed) via likelihood ratio test (Patel et al. 2018). Significant genetic effects (p < 0.05) indicate the significance of heritability estimates. The univariate model can be extended to a bivariate model through Cholesky decomposition (Patel et al. 2018) in order to estimate the genetic correlation (r_g_) and the shared heritability (H^2^) between two phenotypes. Genetic correlation (r_g_) estimates the extent to which the genetic effects that explain the variance in one trait also explain the variance in the second trait. Thus, the order in which the traits are studied is important. For this study, the larger structure (ex: total brain volume) was selected as the leading phenotype. Shared heritability (H^2^) quantifies the degree to which the genetic factors that are shared between the two phenotypes under study are heritable (Patel et al. 2018).

Univariate heritability (H^2^) was estimated in the following measures:

- Total brain volume (TBV), left striatal volume, right striatal volume, left thalamic volume, right thalamic volume, left pallidal volume and right pallidal volume (all volumes adjusted for sex and age) (see Figure 4);
- Left striatal volume, right striatal volume, left thalamic volume, right thalamic volume, left pallidal volume and right pallidal volume (all volumes adjusted for sex, age and TBV) (see Figure S3);
- Sex and age-adjusted vertex-wise measures (surface area and displacement examined separately) in the left and right striatum, thalamus and globus pallidus (see Figure 5);
- Sex, age and TBV-adjusted vertex-wise measures (surface area and displacement examined separately) in the left and right striatum, thalamus and globus pallidus (see Figure S6);

Shared heritability estimates (H^2^) and genetic correlations (r_g_) were computed between TBV and the following measures:

- Left striatal volume, right striatal volume, left thalamic volume, right thalamic volume, left pallidal volume and right pallidal volume (all volumes adjusted for sex and age) (see Figure S5 for shared heritability and Figure S6 for genetic correlation estimates);
- Sex and age-adjusted vertex-wise measures (surface area and displacement examined separately) in the left and right striatum, thalamus and globus pallidus (see Figure S7 for shared heritability and Figure S8 for genetic correlation estimates).

### 4.3 Intraspecies analysis: partial least squares correlation (PLSC) analysis in the Human Connectome Project (HCP) sample

We performed a Partial Least Squares Correlation (PLSC) analysis on the HCP subjects to generate latent variables (LV) relating vertex-wise surface area and displacement in the subcortical structures of interest to behavior (see Figure 1.3.E). The PLSC algorithm applies a singular value decomposition to the covariance matrix between two sets of Z-scored input features: the brain features (vertex-wise surface area and displacement, examined separately in this study and residualized for sex and age) and behavioral features (30 attributes selected from the HCP restricted behavioral and demographic data, residualized for sex and age). Pyls 0.01 (https://github.com/rmarkello/pyls) was used to run the PLSC analysis in Python 3.6.8. Each LV captures what linear combination of brain shape measurements covaries the most with what linear combination of behavioral attributes. The significance of each LV is assessed via permutation testing: a distribution of singular values specific to the LV of interest is generated by shuffling the rows of the brain matrix and performing the SVD decomposition of the new brain-behavior covariance matrix and repeating this 10 000 times. Significance can then be determined via null-hypothesis test. To assess the reliability of the contribution of each vertex to a specific LV, the PLSC algorithm is applied to bootstrapped samples of the dataset 10 000 times in order to obtain a distribution for the loading for each brain variable onto each LV. This allows for the computation of the bootstrap ratio (BSR) of each vertex for each LV, corresponding to that vertice’s loading onto the LV of interest multiplied by the LV’s singular value and divided by the standard error obtained through the bootstrap resampling process. Thus, a high BSR value magnitude indicates a strong and consistent effect across the 10 000 bootstraps.

### 4.4 Spin-test between subcortical structure surface maps

We adapted the Alexander-Bloch cortical surface spin-test (Alexander-Bloch *et al*., 2018) to the striatum, thalamus and globus pallidus in order to assess the significance between the correlations that we found between the brain maps generated in this study (heritability, interspecies differences and LV-specific BSR maps for vertex-wise surface area and displacement in the striatum, thalamus and globus pallidus) (see Figure 1.4). To compare two structure-specific surface maps, this test generates a distribution of the correspondence statistic of interest and a p-value is obtained by calculating the percentage of randomly generated correspondence estimates that are above the observed correspondence statistic. The first surface map is projected onto a sphere, permuted by a random angle and projected back onto the structure of interest in order to generate a correspondence estimate between the permuted surface map and the second untouched surface map. This process is repeated 1000 times in order to generate a distribution for the statistic of interest. For the following study, we adapted the openly available cortical spin test script (https://github.com/spin-test/spin-test) to the subcortex by replacing the original sphere file with 3 spheres of 13000, 6500 and 3000 vertices for the striatum, thalamus and globus pallidus, respectively. This test is implemented bilaterally. We selected Pearson’s correlation coefficient as our correspondence statistic and performed 5% false discovery rate (FDR) correction across 42 comparisons (Benjamin & Hochberg 1995). For bilateral vertex-wise surface area, we used the subcortical spin-test to assess spatial correspondence between the following 5 pairs of surface maps: interspecies differences and heritability, areal LV 1 BSR and interspecies differences, areal LV 2 BSR and interspecies differences, areal LV 1 BSR and heritability and areal LV 2 BSR and heritability. For each of these 5 pairs, the spin-test was conducted on a per structure basis for a total of 15 comparisons (5 pairs x 3 surfaces). For bilateral vertex-wise displacement, we performed a total of 27 comparisons across 9 pairs of surface maps: interspecies differences and heritability, LV 1 BSR and interspecies differences, LV 2 BSR and interspecies differences, LV 3 BSR and interspecies differences, LV 4 BSR and interspecies differences, LV 1 BSR and heritability, LV 2 BSR and heritability, LV 3 BSR and heritability, LV 4 BSR and heritability.

