## Supplemental Figures 1-8 for "Variation in subcortical anatomy: relating interspecies differences, heritability, and brain-behavior relationships"

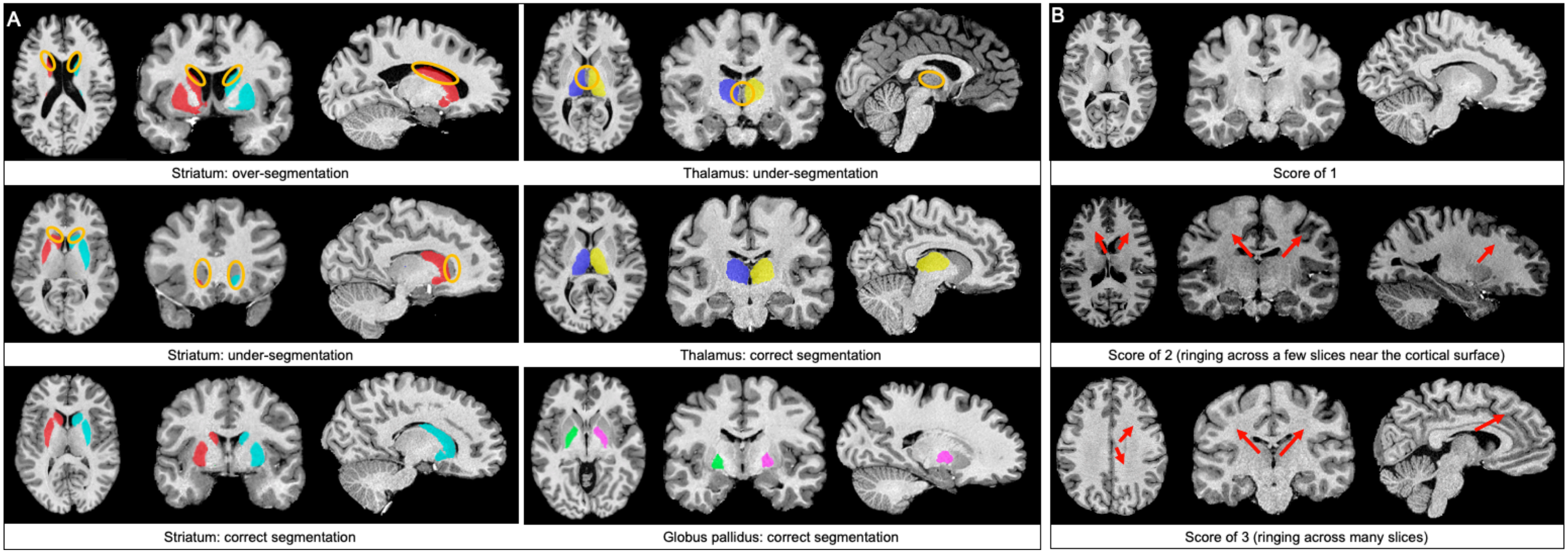
A
Striatum: over-segmentation
Striatum: under-segmentation
Striatum: correct segmentation
Thalamus: under-segmentation
Thalamus: correct segmentation
Globus pallidus: correct segmentation
B
Score of 1
Score of 2 (ringing across a few slices near the cortical surface)
Score of 3 (ringing across many slices)

**Figure S1. Examples of motion quality control ratings and subcortical structure segmentations.**

A) Motion quality control ratings after data preprocessing with minc-bpipe; red arrows point in the direction of motion artifacts (ringing); images in each panel are being displayed along axial (left), coronal (center) and sagittal (right) planes. B) Good and bad segmentations for the striatum, thalamus and globus pallidus; orange circles indicate areas that are over or under segmented; images are being displayed along axial (left), coronal (center) and sagittal (right) planes.

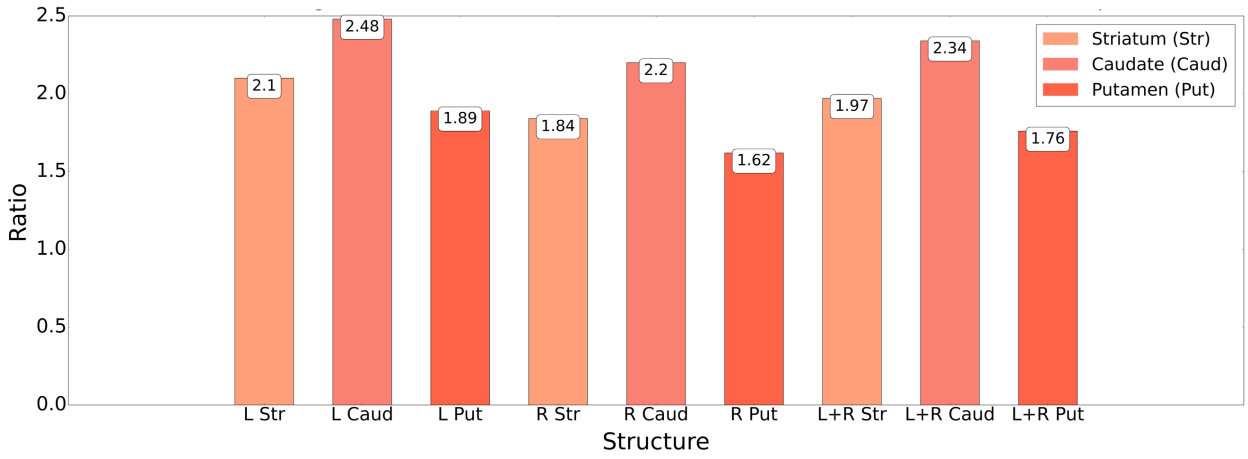


**Figure S2. Ratio of striatum-specific volumes in humans relative to striatum-specific volumes in chimpanzees.**
Show for the left (L), right (R) and bilateral (L+R) striatum (Str), caudate (Caud) and putamen (Put).

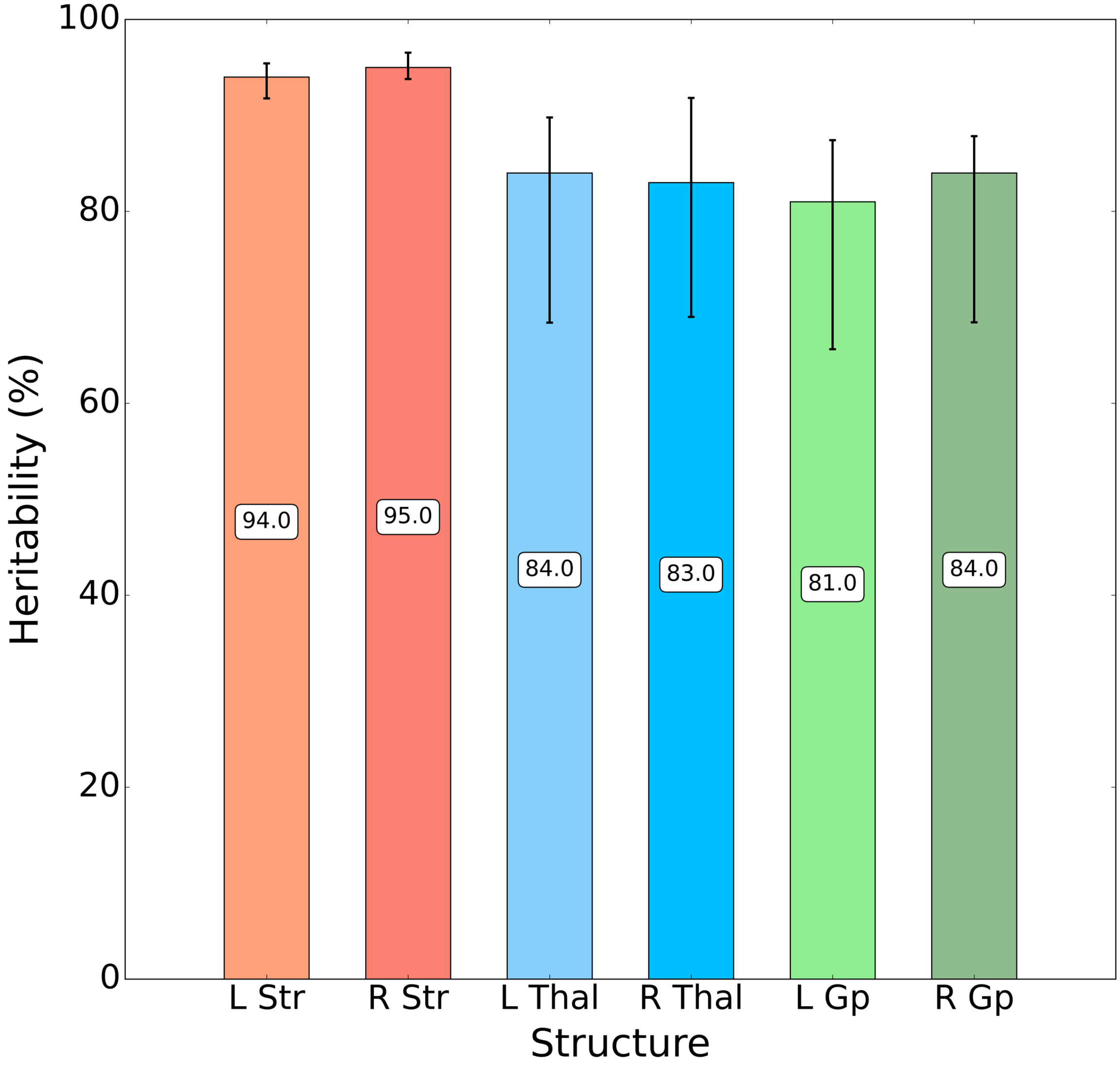


**Figure S3. Heritability of ipsilateral structure-specific volumes, adjusted for sex, age and TBV.**
Shown for the left (L) and right (R) striatum (Str), thalamus (Thal) and globus pallidus (GP). Black bars indicate 95% confidence intervals.

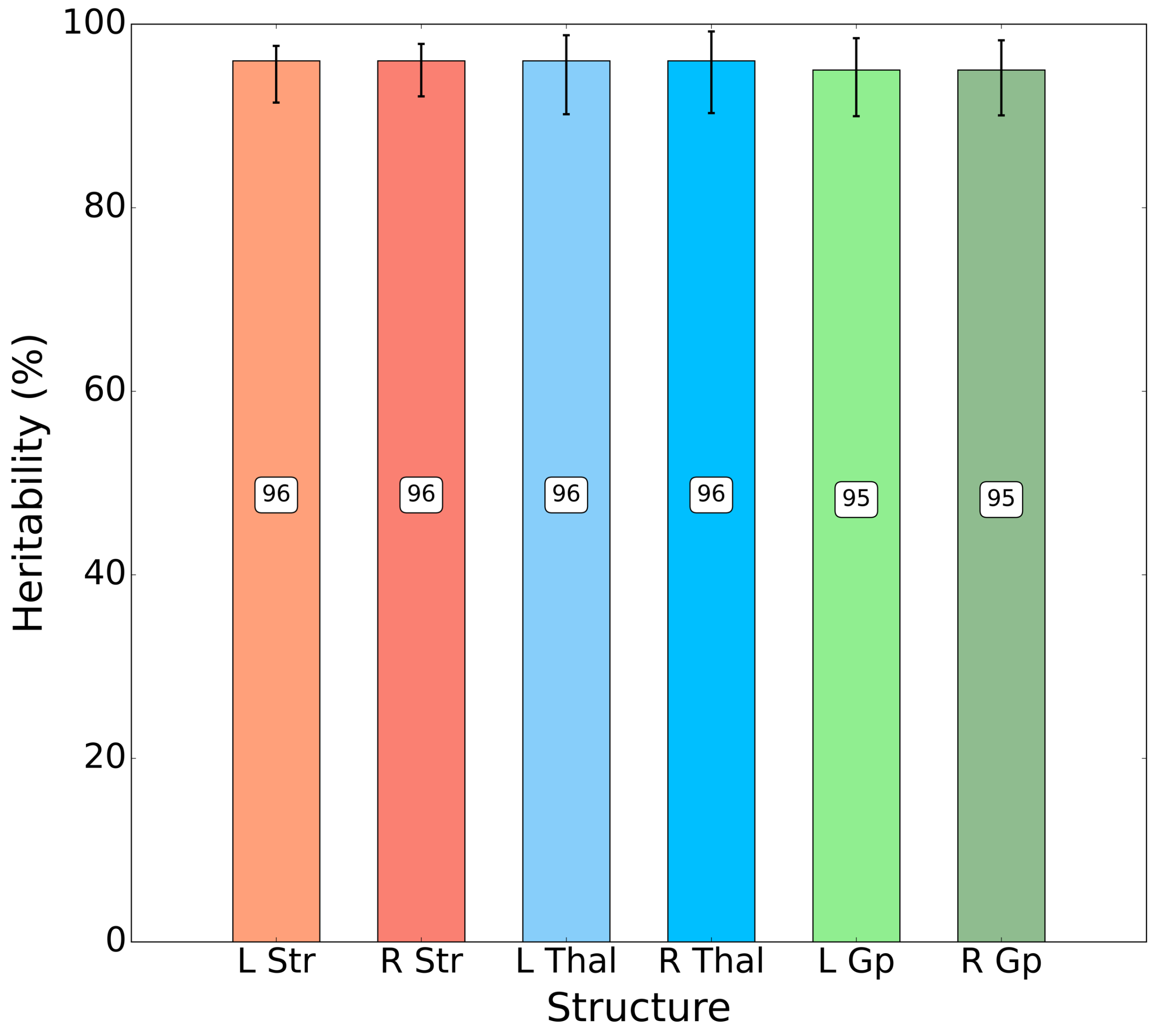


**Figure S4. Shared heritability between TBV and ipsilateral structure-specific volumes, adjusted for sex and age.**
Shown for the left (L) and right (R) striatum (Str), thalamus (Thal) and globus pallidus (GP). Black bars indicate 95% confidence intervals.

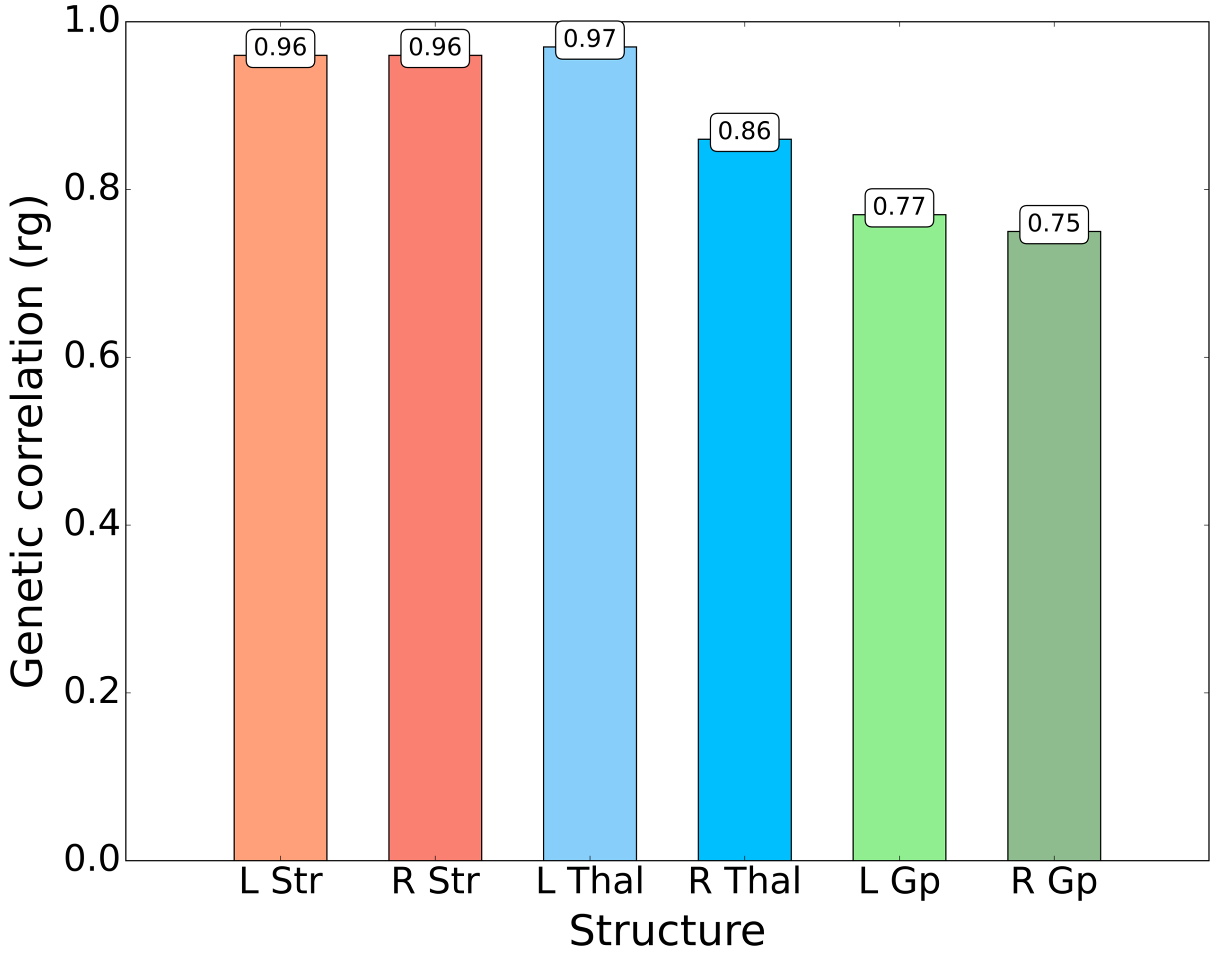


**Figure S5. Genetic correlations (rg) between TBV and ipsilateral structure-specific volumes, adjusted for sex and age.**
Shown for the left (L) and right (R) striatum (Str), thalamus (Thal) and globus pallidus (GP).

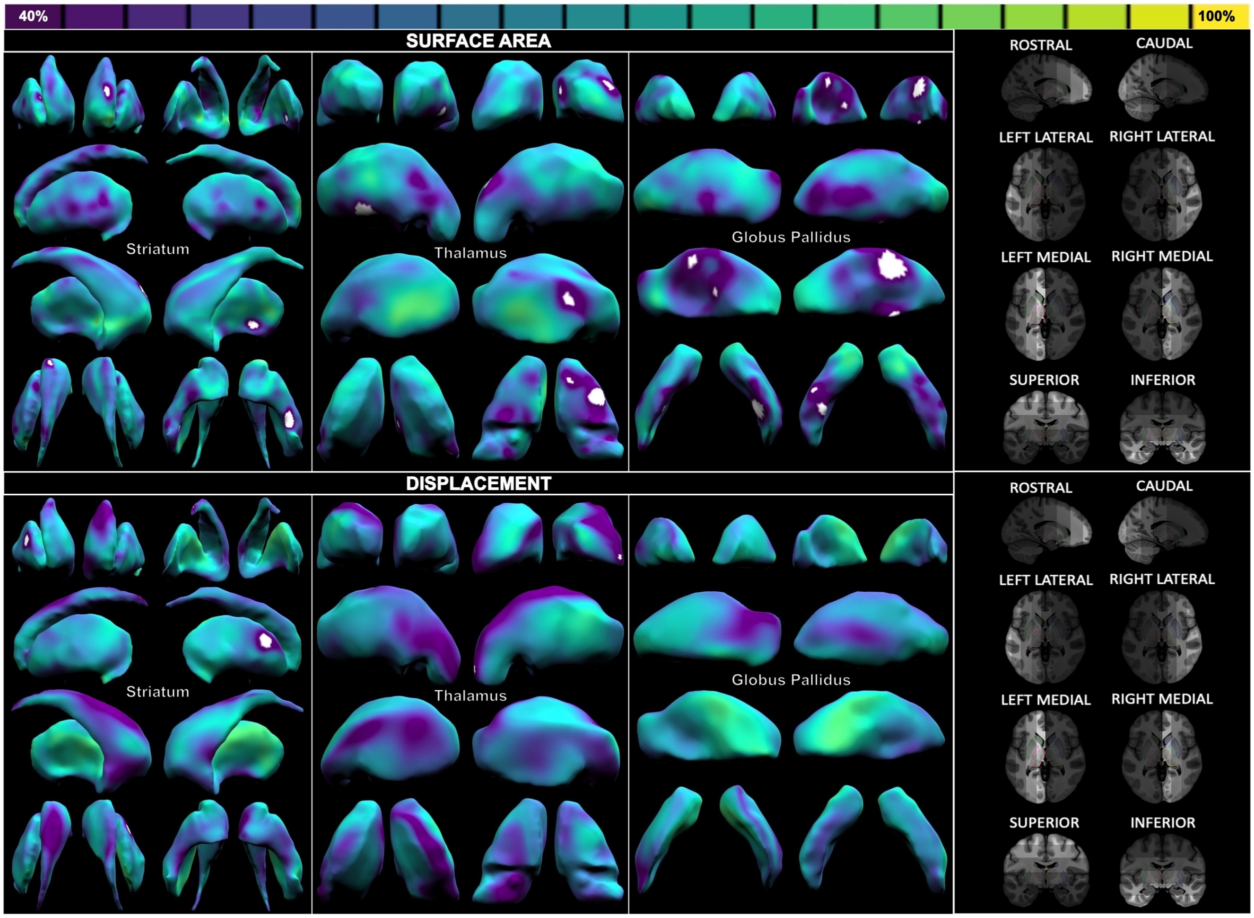


**Figure S6. Heritability of sex, age and TBV-adjusted vertex-wise surface area (top row) and displacement (bottom row) in the striatum, thalamus and globus pallidus.**
The views of the structures on display are shown in the column on the right-hand side; brighter areas on the brain indicate the view from which the structure is being seen. Values in white are not significant (5% FDR correction). Ranges go from 40% (dark purple) to 100% (yellow).

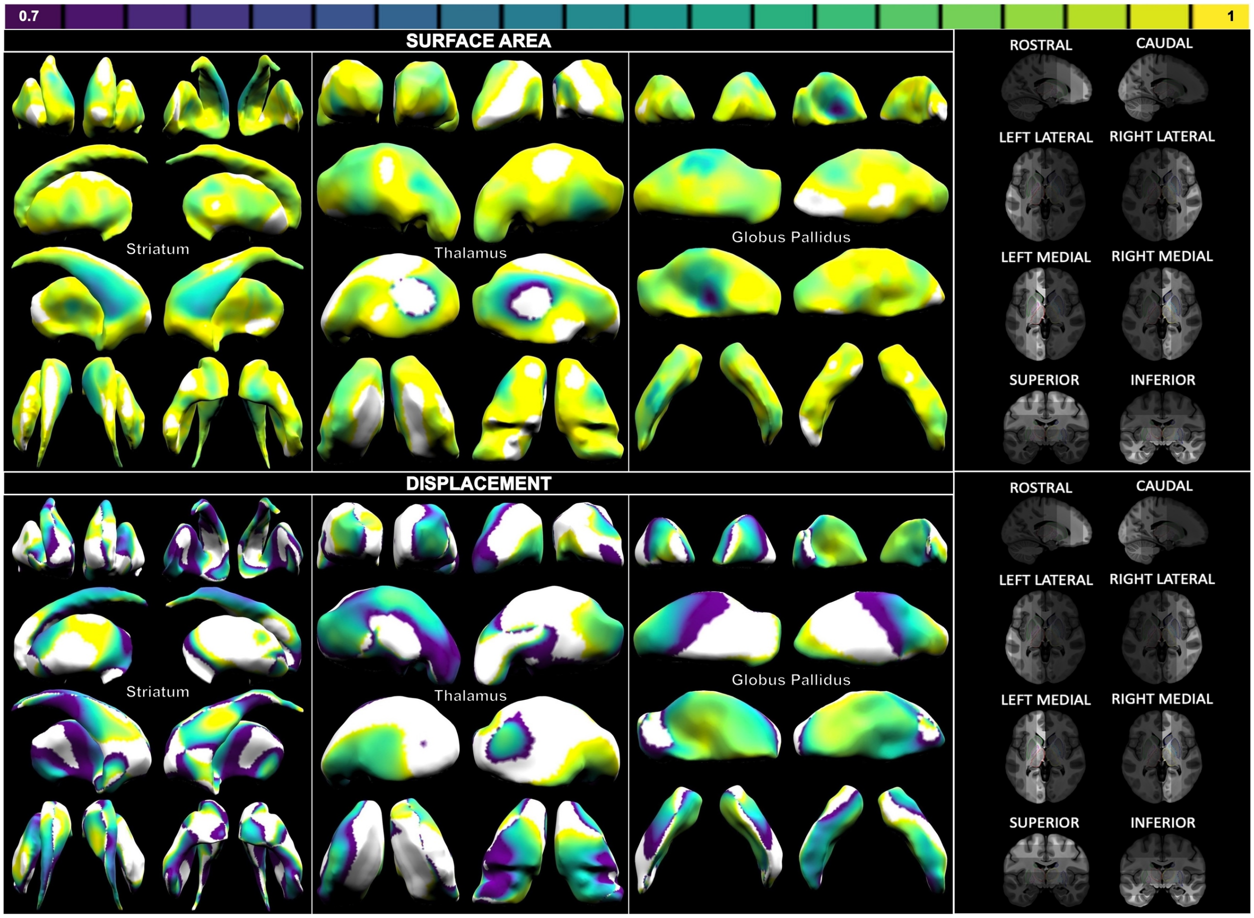


**Figure S7. Shared heritability between TBV and sex and age-adjusted vertex-wise surface area (top row) and displacement (bottom row) in the striatum, thalamus and globus pallidus.**

The views of the structures on display are shown in the column on the right-hand side; brighter areas on the brain indicate the view from which the structure is being seen. Values in white either failed optimization (heritability value below 0% or above 100%) or 5% FDR correction. Ranges go from 70% (dark purple) to 100% (yellow).

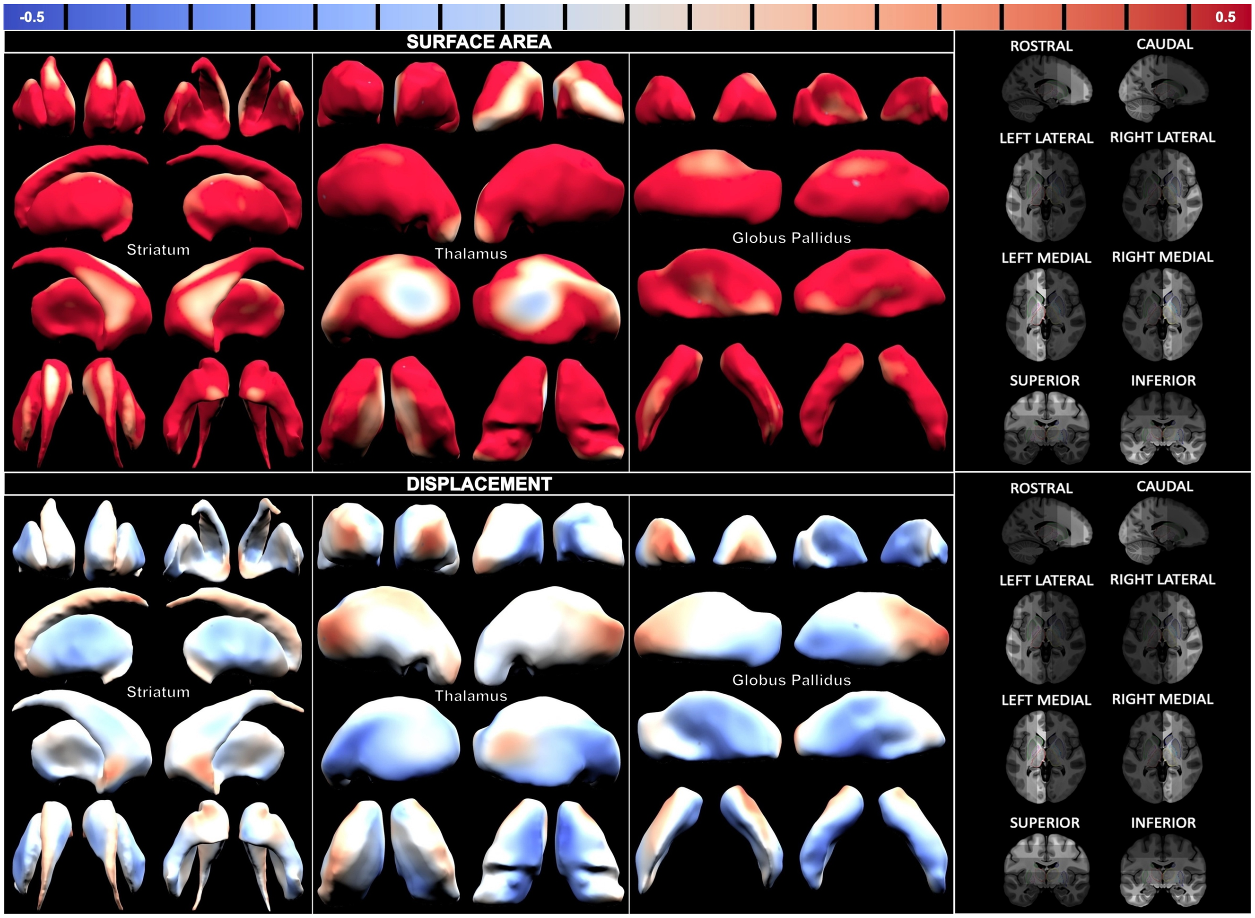


**Figure S8. Genetic correlation (rg) between TBV and sex and age-adjusted vertex-wise surface area (top row) and displacement (bottom row) in the striatum, thalamus and globus pallidus.**
The views of the structures on display are shown in the column on the right-hand side. Ranges go from -0.5 (blue) to 0.5 (red).
